# Th17 cells contribute to combination MEK inhibitor and anti-PD-L1 therapy resistance in *KRAS/p53* mutant lung cancers

**DOI:** 10.1101/2020.06.07.136366

**Authors:** David H. Peng, B. Leticia Rodriguez, Lixia Diao, Pierre-Olivier Gaudreau, Aparna Padhye, Joshua Ochieng Kapere, Caleb A. Class, Jared J. Fradette, Laura Gibson, Limo Chen, Jing Wang, Lauren A. Byers, Don. L Gibbons

**Affiliations:** Department of Thoracic/Head and Neck Medical Oncology, The University of Texas MD Anderson Cancer Center, Houston, TX 77030, USA; Perlmutter Cancer Center, NYU Langone Health, 550 First Avenue, Smilow Building 10^th^ Floor, Suite 1010, New York, NY 10016, USA; Department of Bioinformatics and Computational Biology, The University of Texas MD Anderson Cancer Center, Houston, TX 77030, USA; Thoracic & Upper GI Cancer Research Laboratories, Research Institute of the McGill University Health Centre, Montreal, QC, Canada; The University of Texas MD Anderson Cancer Center UT Health Graduate School of Biomedical Sciences, Houston, TX 77030, USA; Department of Biostatistics, The University of Texas MD Anderson Cancer Center, Houston TX 77030, USA; Department of Molecular and Cellular Oncology, The University of Texas MD Anderson Cancer Center, Houston, TX 77030, USA

**Keywords:** Lung cancer, MEK inhibitor, immunotherapy, Th17, IL-17

## Abstract

Understanding resistance mechanisms to targeted therapies and immune checkpoint blockade in mutant KRAS lung cancers is critical to developing novel combination therapies and improving patient survival. Here, we show that MEK inhibition enhanced PD-L1 expression while PD-L1 blockade upregulated MAPK signaling in mutant KRAS lung tumors. Combined MEK inhibition with anti-PD-L1 synergistically reduced lung tumor growth and metastasis, but tumors eventually developed resistance to sustained combinatorial therapy. Multi-platform profiling revealed that resistant lung tumors have increased infiltration of Th17 cells, which secrete IL-17 and IL-22 cytokines to promote lung cancer cell invasiveness and MEK inhibitor resistance. Antibody depletion of IL-17A in combination with MEK inhibition and PD-L1 blockade markedly reduced therapy-resistance *in vivo*. Clinically, increased expression of Th17-associated genes in melanoma patients treated with PD-1 blockade predicted poorer overall survival and response. Our study validates a triple combinatorial therapeutic strategy to overcome resistance to combined MEK inhibitor and PD-L1 blockade.

## Introduction

Non-small cell lung cancer (NSCLC) is the leading cause of cancer-related deaths due to late-stage disease presentation, metastasis, and resistance to conventional therapies. Approximately 30% of patients with lung adenocarcinoma possess an activating KRAS mutation, which currently lacks approved pharmacological drugs to effectively target the oncogenic protein^1, 2^. Although MEK is a canonical downstream effector of activated mutant KRAS in the MAPK signaling pathway, MEK inhibitors have failed to yield clinical benefit in KRAS mutant cancers^3, 4^. Previous studies have demonstrated that acquisition of common secondary mutations, such as p53, promotes resistance to MEK inhibitors^5^. Additional studies by our group utilizing *Kras;p53* (KP) mutant mouse lung tumor models^6^ demonstrate that epithelial subpopulations of lung cancer cells are responsive to MEK inhibitors, while drug resistant lung cancer cells undergo a ZEB1-dependent epithelial-to-mesenchymal transition (EMT)^7^. Conversely, our prior studies also demonstrate that mesenchymal KP lung tumors are more responsive to PD-L1/PD-1 axis immune checkpoint blockade compared to epithelial KP tumors, due to a ZEB1-mediated upregulation of PD-L1 and other checkpoint proteins in mesenchymal cells^8, 9, 10^.

Although the implementation of PD-1 or PD-L1 immune checkpoint blockade has significantly improved lung cancer patient survival, only a minority of patients show durable response to treatment, suggesting innate or acquired resistance to immunotherapies^11^. Our reported findings suggest that the two distinct subpopulations of lung cancer cells have complementary responses to the individual treatments, providing a potential rationale to combine MEK inhibitors with PD-L1 blockade to overcome resistance to the individual therapies, complementing an on-going clinical trial at MD Anderson (ClinicalTrials.gov Identifier: NCT03225664) implementing a comparable double combination treatment group.

Here, we first show that the combination of MEK inhibition with PD-L1 blockade significantly reduced KP lung tumor growth and metastasis compared to monotherapy treatments. We observed that the initial response to the drug combination was unsustainable with long-term treatment, as primary lung tumors eventually developed resistance. Cytokine array profiling revealed that resistant tumors had increased infiltration of Th17 CD4^+^ T cells, which secrete the tumor-promoting cytokines IL-17 and IL-22^12^. Antibody depletion of IL-17A in combination with MEK inhibition and PD-L1 blockade produced a durable reduction in lung tumor growth, metastasis, and prevented the development of tumor resistance. Gene expression analysis of melanoma patients treated with PD-1 blockade revealed that increasing levels of Th17-associated gene signatures predicted poorer overall survival and response to immune checkpoint blockade. Our findings reveal the molecular rationale for combining MEK inhibitors with PD-L1 blockade, identify the mechanism of combinatorial drug resistance, identify potential predictive markers of immunotherapy response, and validate a promising triple combinatorial treatment strategy for patients with KRAS mutant lung cancer.

## Results

### MEK inhibition increases PD-L1 expression while PD-L1 blockade upregulates MAPK signaling

Previous work from our lab demonstrated that epithelial subpopulations of mutant KRAS lung cancers are responsive to MEK inhibitors while mesenchymal cells within the tumors are resistant^7^. Therefore, we sought to identify potential molecular targets that are specific to mesenchymal subpopulations to synergize with MEK inhibitor treatment. We utilized reverse phase protein array (RPPA) analysis^13, 14^ of heterogeneous syngeneic 344SQ KP lung tumors previously treated with the MEK inhibitor selumetinib (AZD6244)^7^ to identify differentially regulated signaling proteins following MEK inhibition. RPPA profiling revealed a significant (FDR<0.05) upregulation of CD274 (PD-L1) in the 344SQ KP tumors when tumor-bearing mice were treated with AZD6244 (Fig. 1A). Validation of the RPPA data by western blotting confirmed an upregulation of PD-L1 (Fig. 1B). To further test our observation, we analyzed MEK inhibitor-sensitive 393P KP tumors that were previously treated with AZD6244 and harvested at either the point of drug sensitivity (393P AZD-S) or after the development of resistance (393P AZD-R)^7^. Consistent with our prior findings^7, 15^, MEK inhibitor-resistant 393P tumors showed an upregulation of the EMT-associated transcription factor ZEB1 that correlated with increased PD-L1 expression (Fig. 1C). Interestingly, 393P tumors showed an increase in PD-L1 expression even at the point of sensitivity to MEK inhibition (Fig. 1C). To determine if the increase in PD-L1 expression is tumor cell-intrinsic rather than a secondary signal from other cell types present in tumor tissues, we treated murine (393P and 344SQ) and human (H2122, H1299, and H358) lung cancer cell lines *in vitro* with AZD6244 and observed an increase in PD-L1 mRNA and protein expression in lung cancer cells following MEK inhibition (Fig 1D and E; Supplementary Fig. S1A and S1B).

**Figure 1.**
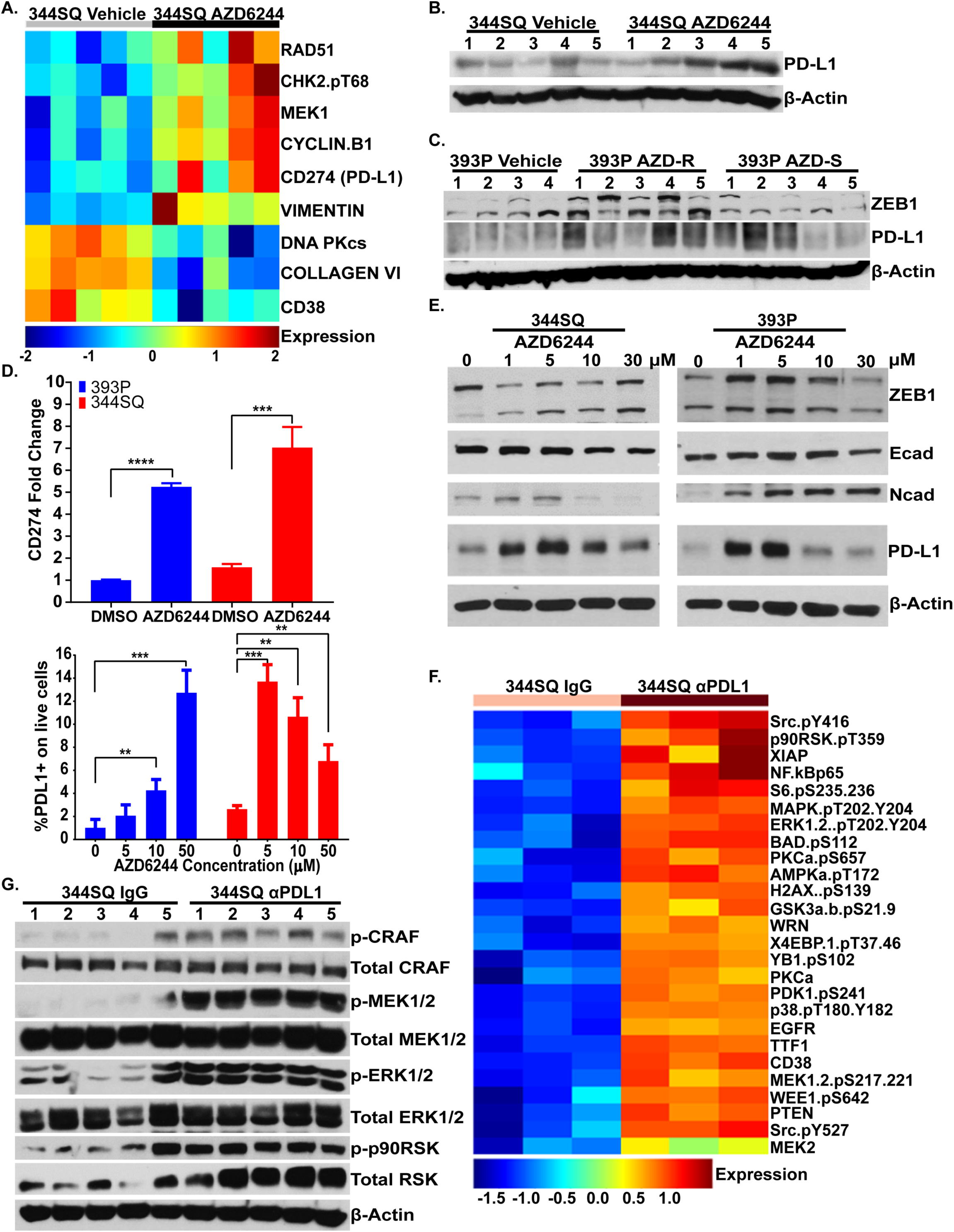
MEK inhibition increases PD-L1 expression while PD-L1 blockade upregulates MAPK signaling. **(A)** Heatmap of reverse phase protein array (RPPA) profile showing statistically significant (FDR<0.05) differentially regulated proteins in 344SQ murine KP syngeneic tumors treated with selumetinib (AZD624) MEK inhibitor or vehicle for 4 weeks. **(B)** Western blot of PD-L1 and β-actin in 344SQ tumors treated with vehicle or AZD6244 for 4 weeks. Number indicates tumor replicate. **(C)** Western blot of ZEB1, PD-L1, and β-actin in 393P tumors treated with AZD6244 for 4 weeks or 8 weeks when tumors were sensitive (AZD-S) or resistant (AZD-R) to MEK inhibition, respectively. Number indicates tumor replicate. **(D)** Top: qPCR analysis of CD274 (PD-L1) expression in 393P and 344SQ cell lines treated with DMSO or 10 µM AZD6244 for 48 hrs. Bottom: Percent of PD-L1+ 393P and 344SQ cells analyzed by flow cytometry following treatment with indicated concentrations of AZD6244 for 48 hrs. **(E)** Western blotting of indicated proteins in 393P and 344SQ cells treated with indicated concentrations of AZD6244 for 48 hrs. **(F)** Heatmap of RPPA profile showing statistically significant (FDR<0.05) differentially regulated proteins in 344SQ tumors treated with PD-L1 blocking antibody or IgG isotype control for 7 weeks. **(G)** Western blotting of indicated proteins of 344SQ tumors treated with IgG isotype control or PD-L1 blocking antibody for 7 weeks. Number indicates tumor replicate.

Previous work by our group demonstrated that mesenchymal KP tumors express high levels of PD-L1 and are responsive to PD-L1 blockade during early phases of tumor growth before developing resistance to immunotherapy^15, 16^. Similarly, to identify upregulated protein targets following PD-L1 blockade resistance, we analyzed RPPA data from 344SQ tumors that previously developed resistance to anti-PD-L1 blockade antibody after 7 weeks of treatment^16^. We observed an increase in the signaling of multiple pathways, including activated phosphorylated MAPK signaling proteins (Fig. 1F), which was validated by western blotting (Fig. 1G). To determine if the increase in MAPK signaling following PD-L1 blockade was due to an immune-mediated response, we performed a co-culture assay with splenocytes and observed an increase in tumor cell MAPK signaling only when 344SQ cells were co-cultured with splenocytes and treated with anti-PD-L1 (Supplementary Fig. S1C). Our findings demonstrate a reciprocal activation of PD-L1 and MAPK signaling when tumors are treated with single-agent MEK inhibitor or PD-L1 blockade, respectively.

### Combination MEK inhibition and PD-L1 blockade initially reduces lung tumor growth and metastasis, but ultimately therapeutic resistance develops

Since MEK inhibition increases PD-L1 expression, while PD-L1 blockade enhances MAPK signaling in lung tumors, we next sought to combine the two treatments to prevent outgrowth of resistant subpopulations from the individual therapies. The combination of AZD6244 with PD-L1 blockade starting at 3 weeks after syngeneic tumor implantation showed a marked synergistic reduction in 344SQ lung tumor growth and metastases that lasted for 8-9 weeks, compared to monotherapy or vehicle/isotype controls (Fig. 2A). However, 344SQ tumors ultimately developed resistance to the combinatorial therapy and by ∼11-12 weeks produced primary tumor size and metastatic disease (Fig. 2A) that were comparable to untreated controls at 9 weeks. Immune profiling of the tumor tissues at the experimental endpoints by flow cytometry showed a significant increase in total CD8^+^ T cells when anti-PD-L1 was administered as a single-agent or in combination with AZD6244 (Fig 2B, Supplementary Fig S2A). Furthermore, memory/effector CD8^+^ T cell subpopulations were significantly increased, while naïve and exhausted CD8^+^ T cells were significantly decreased in the combinatorial treatment group at the point of efficacy (Fig. 2C, Supplementary Fig. S2A). However, despite the transient benefits in the intratumoral T cell populations, exhausted CD8^+^ T cells (PD-1^+^TIM-3^+^) were significantly increased when tumors eventually developed resistance to combination treatment (Fig. 2C).

**Figure 2.**
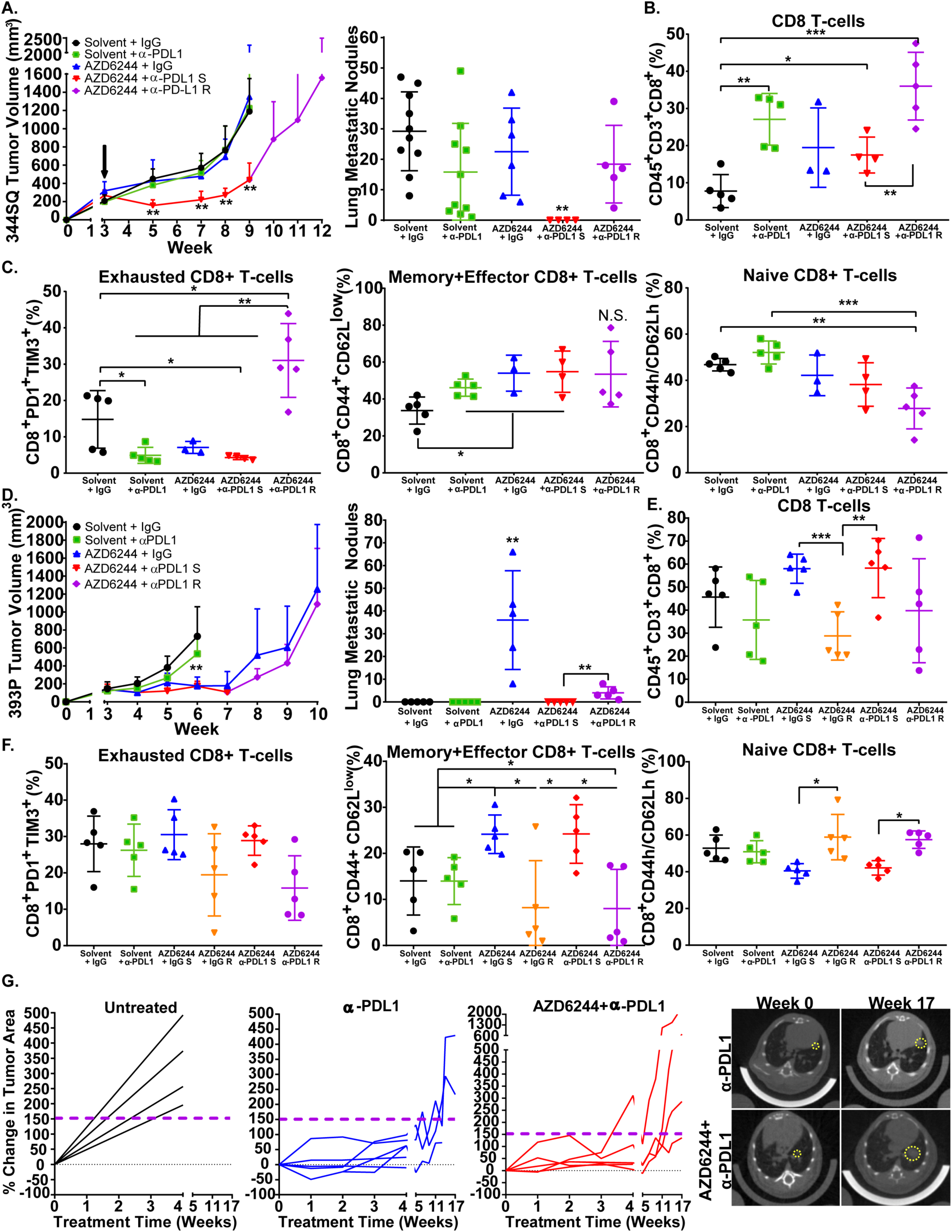
MEK inhibition in combination with PD-L1 blockade initially reduces lung tumor growth and metastasis, but ultimately develops resistance to therapy. **(A)** Left: *In vivo* tumor volume measurements at indicated time points for 344SQ subcutaneous tumors in syngeneic wild-type mice after daily treatment with 25 mg/kg AZD6244 or weekly treatment with 200 µg PD-L1 blocking antibody as single-agents or in combination. Treatment start time denoted by black arrow, sensitive group denoted by (S), resistant group denoted by (R). Right: Quantification of lung metastatic surface nodules in the indicated experimental groups at the endpoint of treatments. **(B)** Percent of CD45+CD3+ total CD8+ T cells in 344SQ tumors with indicated treatment groups at endpoint of experiment in **(A)**. **(C)** Left: Percent of PD1+TIM3+ exhausted CD8+ T cells. Middle: Percent of CD44+CD62L-memory/effector CD8+ T cells. Right: Percent of CD44+CD62+ naïve CD8+ T cells. All populations gated from CD8+ T cells in **(B)**. **(D)** Left: *In vivo* tumor volume measurements at indicated time points for 393P subcutaneous tumors in syngeneic wild-type mice after daily treatment with 25 mg/kg AZD6244 or weekly treatment with 200 µg PD-L1 blocking antibody as single-agents or in combination. Treatment start time denoted by black arrow. Treatment start time denoted by black arrow, sensitive group denoted by (S), resistant group denoted by (R). Right: Quantification of lung metastatic surface nodules in the indicated experimental groups at the endpoint of treatments. **(E)** Percent of CD45+CD3+ total CD8+ T cells in 393P tumors with indicated treatment groups at endpoint of experiment in **(D)**. **(F)** Left: Percent of PD1+TIM3+ exhausted CD8+ T cells. Middle: Percent of CD44+CD62L-memory/effector CD8+ T cells. Right: Percent of CD44+CD62+ naïve CD8+ T cells. All populations gated from CD8+ T cells in **(E)**. **(G)** Left: Percent change in overall lung tumor area of age-matched *Kras*^*G12D*^;*p53*^*-/-*^ (KP) at indicated time points following weekly treatment with 200 µg PD-L1 blocking antibody monotherapy or in combination with daily treatments of 25 mg/kg AZD6244, as assessed by micro-CT imaging of mouse lungs. Right: Representative cross-sectional micro-CT images of KP mouse lungs before indicated treatment (Week 0) and treatment endpoint (Week 17). Yellow circles outline representative target lesions.

Similarly, we treated MEK inhibitor-sensitive, epithelial, 393P KP tumors with the combinatorial therapy and observed no significant difference in primary tumor growth rate between single-agent AZD6244 and the combination group (Fig. 2D). Consistent with our prior publication, the epithelial 393P tumors also developed resistance to either the single-agent or combination treatments, but interestingly, the combination group showed a reduction in metastases compared to the AZD6244 monotherapy resistant group, even at the point (10 weeks) when the primary tumors displayed resistance (Fig. 2D). Immune profiling of 393P tumor tissues at the experimental endpoints or after 4 weeks of treatment showed an increase in total CD8+ T cells only when tumors were treated with AZD6244 monotherapy or in combination with anti-PD-L1 (Fig. 2E and Supplementary Fig. S2B). Again, we only observed a significant increase in memory/effector CD8^+^ T cell subpopulations during the responsive period to treatment with AZD6244 or combination therapy (Fig. 2F and Supplementary Fig. S2B). Nanostring analysis on immune markers in 344SQ and 393P tumors in the different treatment groups confirmed our flow cytometry data, showing that treatment of tumors resulted in increase of T cell gene signatures. Notably, tumors that were resistant to combination treatment showed a decrease in T cell gene signatures (Supplementary Fig. S2C and S2D). Analysis of additional immune cell populations, including CD4^+^ T cell subgroups and antigen presenting cells (APC), in 344SQ and 393P tumors after 4 weeks of treatment did not show statistically significant or consistent changes between the treatment groups (Supplementary Fig. S2C-H).

Our prior work demonstrated that autochthonous mutant *Kras*^*G12D*^;*p53*^*-/-*^ (KP) lung tumors in the genetically engineered mouse model (GEMM) were resistant to AZD6244 monotherapy^7^, but were partially responsive to anti-PD-L1 blockade for ∼12 weeks before developing resistance^16^. To test out findings from syngeneic tumor models in GEMMs, we treated KP mice with anti-PD-L1 alone or in combination with AZD6244 to determine if combination therapy could prevent resistant tumor outgrowth. Micro-CT imaging of mice lungs showed an initial reduction in lung tumor growth in mice that received the combination therapy or anti-PD-L1 monotherapy when compared to untreated controls (Fig. 2G). However, lung tumors developed resistance to single-agent and combination therapy after 17 weeks of treatment, consistent with our prior reports (Fig. 2G)^16^.

### Combination therapy resistant syngeneic and autochthonous tumors have increased levels of Th17 CD4^**+**^ **T cells**

Although combinatorial therapy in 344SQ and 393P tumors exhibited differences in tumor growth and metastatic phenotypes, both tumor types exhibited similar changes in CD8^+^ T cell patterns at the point of sensitivity and resistance to the respective drug treatments, suggesting a mutual secondary immune cell infiltrate following CD8^+^ T cell infiltration that promotes resistance. To identify specific immune cell populations and cytokines upregulated in drug resistant 393P and 344SQ tumors, we performed a qPCR array analyzing multiple cytokines, immune-associated receptors and transcription factors. Data from the immune qPCR array of 344SQ and 393P tumors treated with individual or combinatorial therapies described in Figure 2A and 2B showed a consistent upregulation of genes associated with Th17 CD4^+^ T cells (IL23, IL17, IL22, and RORγt) in resistant 393P and 344SQ tumors (Fig. 3A, B; Supplementary Fig. S3A and S3B). Th17 cells are a subset of pro-inflammatory CD4^+^ T cells that express RORγt and secrete the tumor-promoting cytokines IL-17 and IL-22^12, 17, 18, 19, 20, 21, 22, 23, 24, 25^. Nanostring analysis of 344SQ tumors that were responsive and resistant to combinatorial treatment confirmed our qPCR array, showing a significant upregulation of IL-17F in resistant 344SQ tumors (Supplementary Fig. S3C). Flow cytometry analysis of 344SQ and 393P tumor tissues showed an increase in CCR5^+^, CCR6^+^, and IL-17^+^RORγt^+^ CD4^+^ T cells in tumors that developed resistance to therapy (Fig. 2C, D; Supplementary Fig. S3D). IHC stains for RORγt in autochthonous KP lung tumors treated with anti-PD-L1 or combination therapy from Figure 2G showed an increase in RORγt^+^ tumor infiltrating cells in KP tumors resistant to either treatment (Fig. 3E).

**Figure 3.**
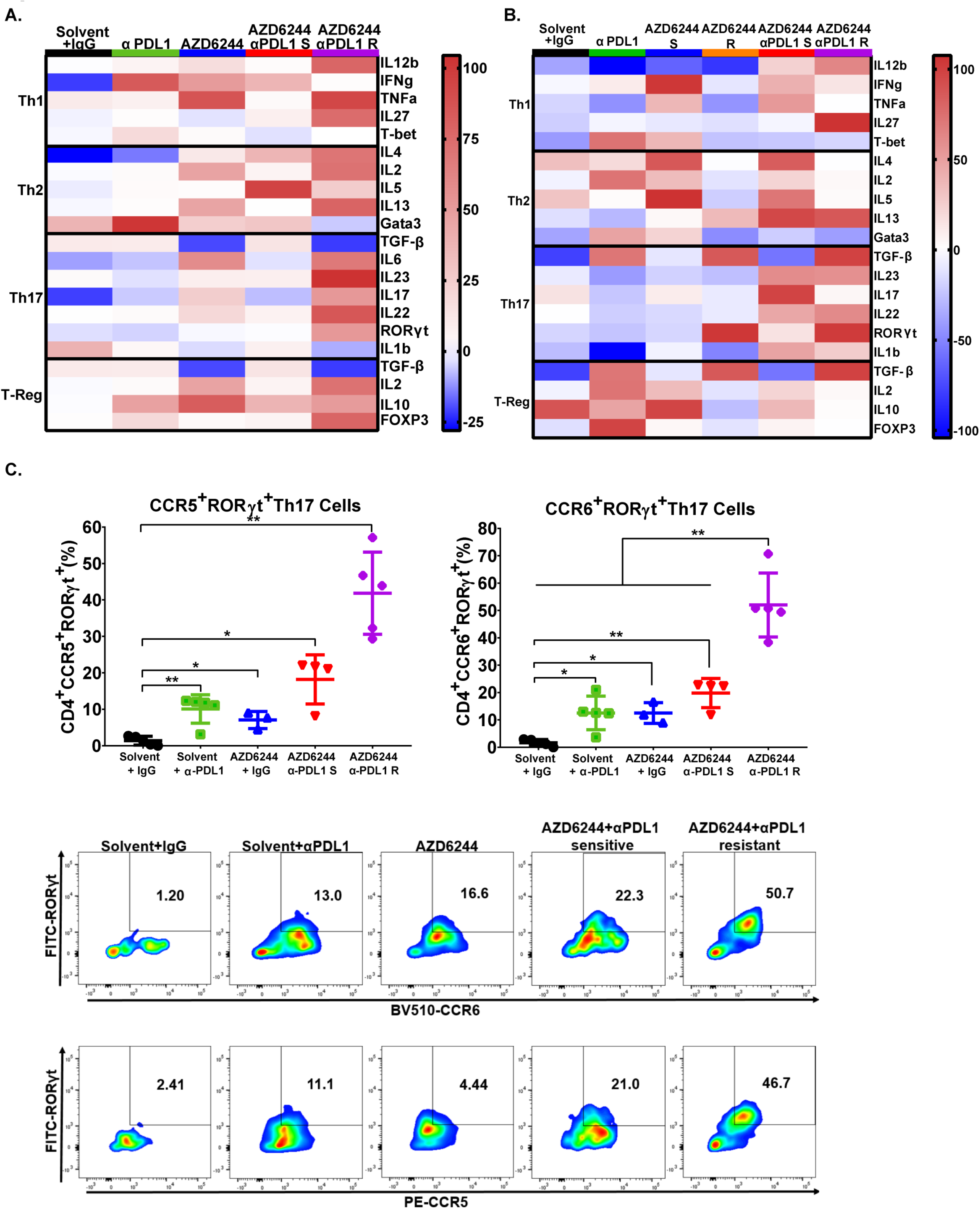
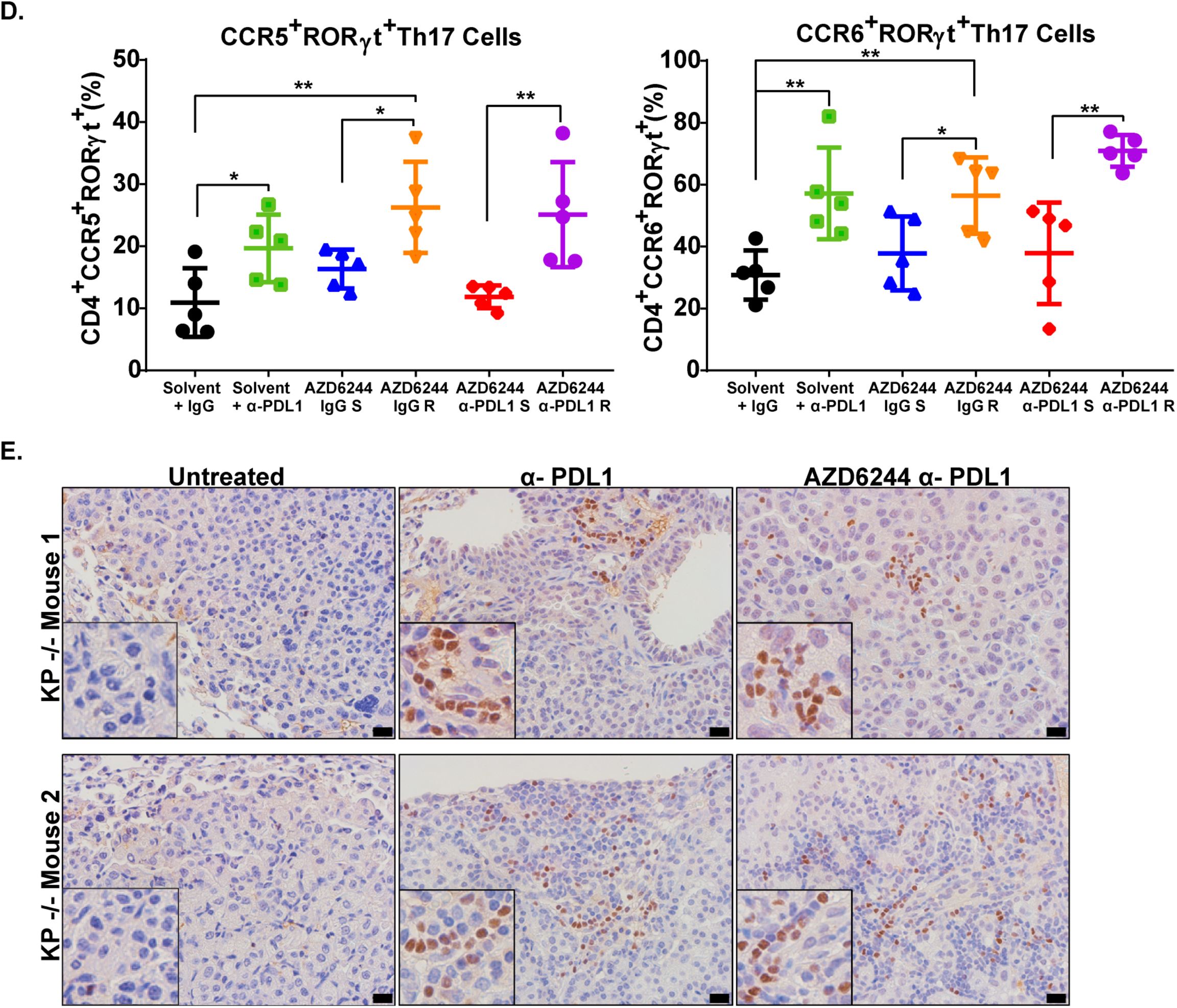
Combination therapy resistant tumors have increased levels of Th17 CD4^+^ T cells. **(A)** Cytokine qPCR array heatmap of 344SQ tumors from the experiment in **Figure 2A**. **(B)** Cytokine qPCR array heatmap of 393P tumors from the experiment in **Figure 2D**. **(C)** Top: Percent of CCR5+RORγt+ and CCR6+RORγt+ Th17 CD4+ T cells in 344SQ tumors from treatment experiment in **Figure 2A**. All populations gated from total CD4+ T cells in **Supplementary Figure S3D**. Bottom: Gating sheme for CCR5+RORγt+ and CCR6+RORγt+ Th17 CD4+ T cells. **(D)** Percent of CCR5+RORγt+ and CCR6+RORγt+ Th17 CD4+ T cells in 393P tumors from treatment experiment in **Figure 2D**. **(E)** RORγt IHC stains of KP mice lung tumors treated weekly with 200 µg of PD-L1 blocking antibody alone or in combination with 25 mg/kg daily treatments of AZD6244 for 17 weeks from the experiment in **Figure 2G**.

### MEK inhibition contributes to differentiation of Th17 cells, which secrete IL-17 and IL-22, promoting tumor cell drug resistance and invasiveness

Since Th17 cells typically require TGF-β, IL-6, and IL-23 stimulation for differentiation^26^, we next wanted to determine if MEK inhibition of lung cancer cells contributed to Th17 cell differentiation. First, we treated murine KP lung cancer cells with AZD6244, and observed an increase in TGF-β, IL-6, and IL-23 expression following MEK inhibition (Fig. 4A). A similar trend was found in human NSCLC cell lines H358 and H1299 treated *in vitro* with AZD6244 (Supplementary Fig. 4A). Additionally, AZD6244 treatment of human lung cancer cells showed a consistent upregulation of Th17-associated genes (TGF-β, IL6, IL23, IL17, IL22, IL1b, and RORγt) (Supplementary Fig. S4A). Next, we co-cultured 393P and 344SQ KP cells with anti-CD3/CD28 activated splenocytes, treated the co-culture with AZD6244, and analyzed splenocyte populations for Th17 cells. *In vitro* co-culture assays showed an increase in CCR5^+^RORγt^+^CD4^+^ T cells with no significant changes in total CD4^+^ T cells when 393P or 344SQ cells were treated with AZD6244 (Fig. 4B). To determine if Th17 cells from co-cultures were secreting IL-17, we co-cultured 344SQ cells with anti-CD3/CD28 activated splenocytes, treated cells with AZD6244, anti-PD-L1, or both in combination, and analyzed secreted IL-17 levels in the conditioned media by ELISA. Treatment of 344SQ cells alone did not show a significant change in secreted IL-17 regardless of treatment (Fig. 4C). However, co-culture of splenocytes with 344SQ cells showed detectable levels of IL-17 in the conditioned media, which was further increased (approximately two-fold) when cells were treated with the combination therapy (Fig. 4C). Next, we tested whether IL-17 and IL-22 cytokines secreted by Th17 cells promoted lung cancer cell drug resistance and invasion. 393P and 344SQ cells treated with recombinant IL-17 and IL-22 were more invasive through Matrigel-coated transwell inserts (Fig. 4D) and became less responsive to *in vitro* AZD6244 MEK inhibition (Fig. 4E).

**Figure 4.**
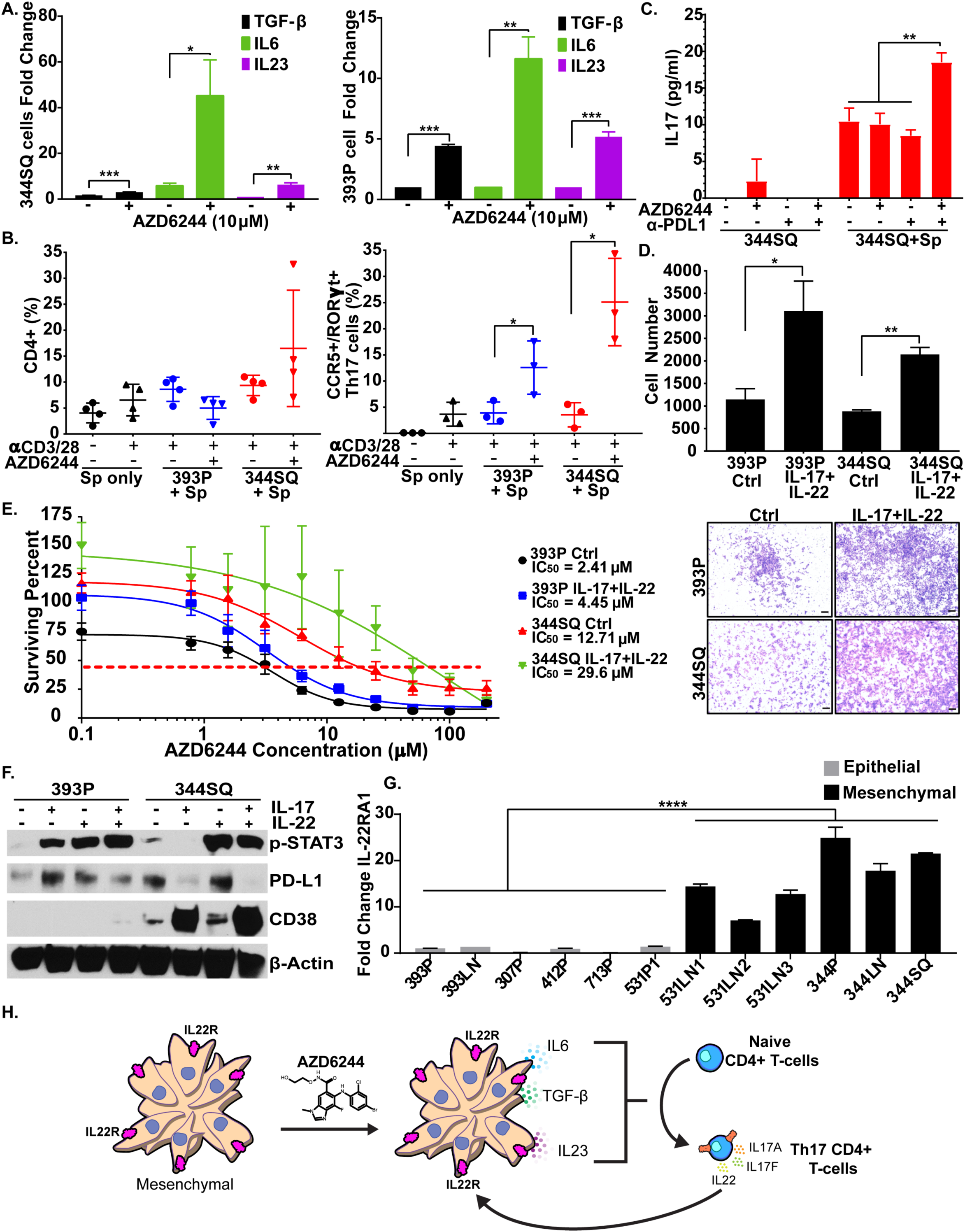
MEK inhibition contributes to differentiation of Th17 cells, which secrete IL-17 and IL-22 to promote drug resistance and invasiveness in lung cancer cells. **(A)** QPCR analysis of TGF-β, IL-6, and IL-23 gene expression in 344SQ (left) and 393P (right) murine KP cell lines after treatment with 10 µM AZD6244 for 48 hrs. **(B)** Left: Percent of CD4^+^ T cells gated from CD45^+^CD3^+^ cells in total anti-CD3/CD28 activated splenocytes (Sp) co-cultured with 393P or 344SQ cells and treated with 10 µM AZD6244 for 96 hrs. Right: Percent of CCR5^+^RORγt^+^ Th17 cells gated from total CD4^+^ T cell population (Left) following co-culture and treatment conditions previously described. **(C)** IL-17 concentration as measured by ELISA in conditioned media of 344SQ cells cultured alone or in co-culture with splenocytes, treated with 10 µM AZD6244 and/or 20 µg/ml PD-L1 blocking antibody for 96 hrs. **(D)** Quantification and representative images of 393P and 344SQ cell invasion through Matrigel-coated transwell inserts ± IL-17 and IL-22 cytokine stimulation. **(E)** *In vitro* cell survival response after 72-hour selumetinib (AZD6244) treatment in 393P and 344SQ cells ± IL-17 and IL-22 cytokine stimulation. **(F)** Westerns blot of indicated proteins in 393P and 344SQ cells treated with IL-17 and/or IL-22 individually or in combination for 96 hrs. **(G)** QPCR analysis of IL22RA1 expression in panel of murine epithelial cells or mesenchymal cells. **(H)** Proposed working model. Initial response to combination therapy promotes release of IL6, TGF-β, and IL23 cytokines by resistant tumor cells. The released cytokines promote the CD4+ T cell differentiation into Th17 cells, which secrete IL-17 and IL22 to promote resistance to MEK inhibition and PD-L1 blockade.

Although Th17-associated cytokines promoted MEK inhibitor resistance and cancer cell invasion in 393P and 344SQ cells, 344SQ tumors were more metastatic compared to 393P tumors after acquiring resistance to combination therapy, suggesting differential response to anti-PD-L1. To that end, western blot analysis of immunosuppressive molecules showed that 393P cells upregulated PD-L1 protein levels when cells were treated with IL-17 or IL-22. Conversely, although 344SQ cells had higher basal levels of PD-L1 as previously described^8^, stimulation of 344SQ cells with IL-17 decreased PD-L1 expression but dramatically increased CD38, which promotes an immunosuppressive tumor microenvironment and resistance to PD-(L)1 immune checkpoint blockade^16^. Additionally, western blots also showed an increase in p-Stat3 when 393P cells were treated with IL-17 or IL-22 individually or in combination, while 344SQ cells had higher basal levels of p-Stat3 and increased p-Stat3 signaling only with IL-22 stimulation (Fig. 4F).

To address the differential response to IL-17 and IL-22 stimulation between 393P and 344SQ cells, we analyzed expression of IL-17 and IL-22 receptors in a panel of mouse KP cell lines. Although IL17RA and Il17RC expression was similar across all cell lines (Supplementary Fig. 4B), we observed a marked increase in IL22RA1 for all metastatic, mesenchymal KP cells (Fig. 4G) compared to the non-metastatic, epithelial cells. In addition, levels of IL22RA1 were elevated in 393P cells following ZEB1 overexpression or acquisition of resistance to AZD6244 (Supplementary Fig. 4C). Conversely, overexpression of miR-200 in mesenchymal 344SQ cells significantly reduced IL22RA1 expression, indicating a strong correlation between EMT and IL22RA1 (Supplementary Fig. 4C). IL22RA1 could also be induced in 393P cells following stimulation with recombinant IL-17A (Supplementary Fig. 4C). Our findings suggest a working model in which MEK inhibition in lung cancer cells induces secretion of Th17 differentiating cytokines IL6, TGF-β, and IL23 (Fig. 4H). Infiltrating Th17 cells then secrete IL-17 and IL-22 to promote primary tumor resistance to combination therapy (Fig. 4H).

### Th17 gene signatures predicts response and overall survival in cancer patients treated with PD-1 blockade therapy

To evaluate the clinical relevance of our findings, we analyzed Th17-associated gene signatures in RNA-seq datasets from patients treated with anti-PD-1 to predict patient response to immune checkpoint blockade therapies. Because datasets for lung cancer patients treated with immunotherapies and with matched pre- and post-treatment samples are not available, we analyzed a publically available, published dataset from melanoma patients treated with anti-PD-1 therapies^27^. Expression analysis of Th17-associated genes (IL17RA, IL17RC, IL22RA1, RORC, CCR5, CCR6, IL6, IL23, TGFB1) in pre-treatment melanoma patient samples revealed that upregulation of IL17RC mRNA correlated with progressive or stable disease, while lower IL17RC mRNA correlated with partial or complete response to anti-PD-1 (Fig. 5A). Gene expression comparisons between pre- and on-treatment melanoma patient samples showed a significant overall increase in IL17RA, TGFB1, and CCR5 when patients received anti-PD-1 (Fig. 5B). Because patients showed variable changes in gene expression for multiple markers, analysis of net changes (directional change in expression between pre- and on-treatment biopsies) in Th17-associated gene expression predicted significant improvement in overall survival in patients who showed decreased expression in RORγt, IL17RA, TGFB1, and CCR5 (Fig. 5C). There was no significant differences in overall survival in patients with net changes in other genes associated with CD4 subpopulations (Supplementary Fig. 5A). Our analyses of clinical data corroborate our experimental findings and identify Th17-related genes as promising markers to predict patient outcome following immune checkpoint blockade.

**Figure 5.**
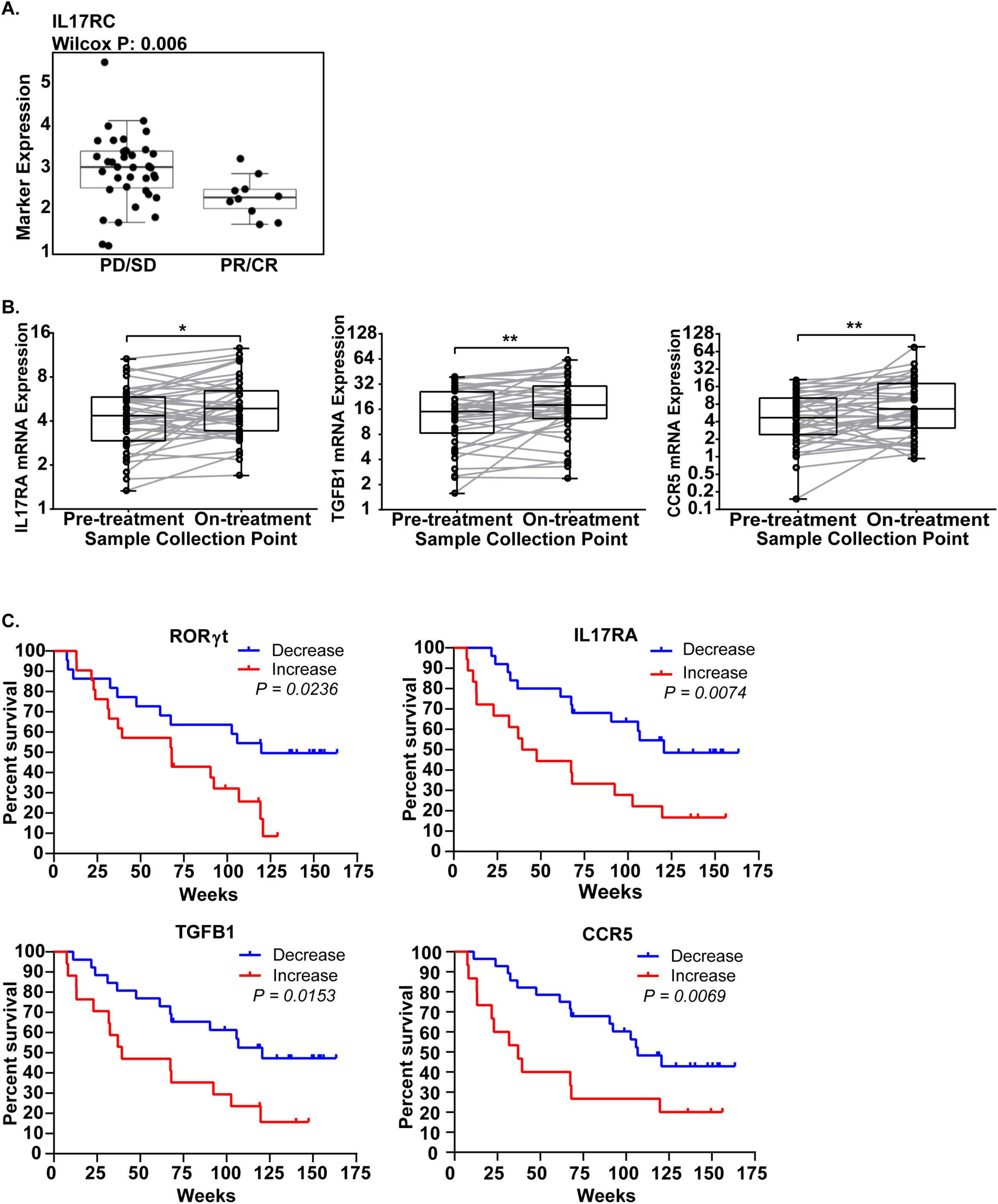
Th17 gene signatures correlate with decrease in survival in IO treated melanoma patients. **(A)** Pre-treatment IL17RC mRNA levels in melanoma patients that exhibited progressive or stable disease (PD/SD) versus partial or complete response (PR/CR) to anti-PD-1 therapy (nivolumab). **(B)** Left: IL17RA, Middle: TGFb-1, and Right: CCR5 mRNA expression levels in melanoma patient samples in pre- and on-treatment of nivolumab. Statistical difference was determined using Wilcoxon matched pair rank test. **(C)** Kaplan-Meier curves predicting survival of nivolumab-treated melanoma patients based on net changes in RORγt, IL17RA, TGFB1, and CCR5. Statistical difference was determined using Log-rank Cox test.

### Neutralizing IL-17 abrogates resistance to combination MEK inhibition and PD-L1 blockade therapy

To test the *in vivo* effects of the increased IL-17 production, therapeutic antibody neutralization of IL-17 was combined with MEK inhibition and PD-L1 checkpoint blockade in 344SQ tumors starting three weeks post-implantation in mice. The triple therapy significantly reduced primary tumor growth and abrogated resistant tumor outgrowth and metastasis compared to tumors treated with the double combination therapy (Fig. 6A). Immune profiling of the tumors at the endpoint of the experiment showed an increase in total CD8^+^ and memory/effector CD8^+^ T cells followed by a decrease in exhausted and naïve CD8^+^ T cells (Fig. 6B). Although total CD4^+^ T cell levels did not change (Supplementary Fig. 5B), we observed a consistent increase in CCR6^+^ and CCR5^+^RORγt^+^ Th17 cells in tumors resistant to the double combination treatment, while tumors treated with the triple combination therapy prevented the increase in Th17 populations (Fig. 6C). Our findings suggest a working model in which single-agent treatment selects for either a resistant epithelial or mesenchymal population, while combinatorial therapy of MEK inhibitor with PD-L1 blockade reduces both epithelial and mesenchymal subpopulations to suppress overall lung tumor growth (Fig. 6D). Although infiltration of IL-17-secreting Th17 cells promotes resistance to the combination therapy, neutralization of IL-17 abrogates resistance, produces sustained activated CD8^+^ T cell levels and lung tumor reduction, validating a promising triple combinatorial treatment strategy to overcome single-agent or combination therapy drug resistance in KRAS driven lung cancers (Fig. 6D).

**Figure 6.**
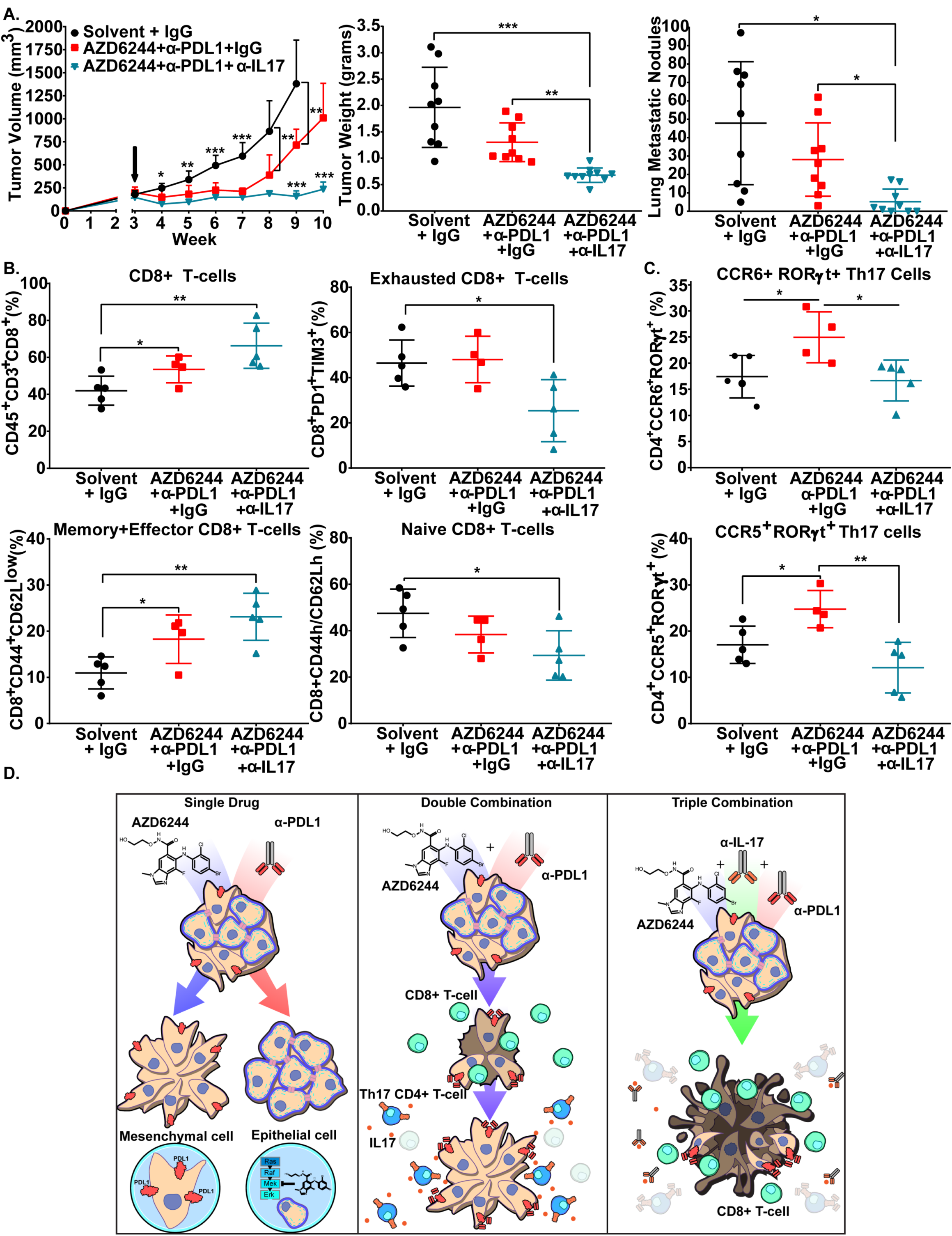
Neutralizing IL-17 abrogates resistance to MEK inhibition and PD-L1 blockade combination therapy. **(A)** Left: *In vivo* tumor volume measurements at indicated time points for 344SQ subcutaneous tumors in syngeneic wild-type mice after daily treatments with 25 mg/kg AZD6244 in combination with weekly treatments with 200 µg PD-L1 blocking antibody. For triple combination group, IL-17A blocking antibody was administered weekly at 200 µg per mouse. Black arrow indicates treatment start time. Middle: Quantification of final 344SQ tumor weight at indicated experimental treatment groups. Right: Quantification of lung metastatic nodules at experimental endpoint. **(B)** Percent of CD45^+^CD3^+^ total CD8^+^ T cells and percentage of exhausted, memory/effector, and naïve CD8^+^ T cell subpopulations gated by the indicated markers. **(C)** Percent of CCR6^+^ and CCR5^+^RORγt^+^ Th17 cells gated from total CD4^+^ T cell population. **(D)** Proposed working model of MEK inhibitor and anti-PD-L1 combinatorial drug resistance. Initial response to combination therapy promotes CD4^+^ T cell differentiation into Th17 cells, which secrete IL-17 and IL-22 to promote resistance to MEK inhibition and PD-L1 blockade. Neutralization of IL-17 in combination with MEK inhibition and PD-L1 blockade abrogates resistant tumor outgrowth.

## Discussion

The failure of MEK inhibitors against KRAS mutant lung cancers and the low percentage of sustained durable responses with anti-PD-1/PD-L1 immune checkpoint blockade therapies emphasizes the need to understand drug resistance mechanisms to improve patient survival. Here we identified a reciprocal expression between PD-L1 and MAPK signaling when lung tumors were treated with MEK inhibitor or anti-PD-L1, respectively, validating a promising dual combinatorial therapy approach previously reported in multiple cancer types^28, 29, 30, 31, 32^. Consistent with previous work^30, 33^, MAPK inhibition in combination with PD-L1 blockade demonstrated an initial reduction in KP lung tumor growth. However, despite the initial response to the double combination therapy and even anti-PD-L1 monotherapy in our KP pre-clinical GEMMs, lung tumors eventually developed resistance to combination therapy. Furthermore, our syngeneic KP tumor models demonstrated that distinct subsets of lung cancer cells with varying baseline sensitivities to MEK inhibitor also developed resistance to double combination therapy, suggesting that a mutual factor promoted resistance. Here, we identified a dramatic increase in Th17 CD4^+^ T cells in resistant tumors in both the syngeneic and KP GEM models. The increase in subpopulations of CD4^+^ T cells is partially corroborated in melanoma phase 1b studies involving MAPK inhibitors and PD-L1 checkpoint blockade combination therapies^29^. Additionally, our data demonstrated that MEK inhibition or PD-L1 blockade alone was sufficient to significantly increase Th17 tumor infiltration. We further demonstrate that IL-17 and IL-22 cytokines secreted by Th17 cells are directly responsible for promoting drug resistance, invasion, and metastasis. Analysis of immune suppressive markers between different lung cancer models showed differential response to Th17 cytokines. In 393P cells, which express low basal levels of PD-L1, IL-17 and IL-22 stimulation upregulated PD-L1 expression, while in 344SQ cells, which express high basal levels of PD-L1 and IL-17, specifically suppressed PD-L1 expression but dramatically upregulated CD38 levels. We previously demonstrated that CD38 is crucial for immune evasion and PD-L1 blockade resistance^16^. Here, we demonstrate that infiltration of Th17 cells and stimulation of cancer cells with IL-17 is a mechanism of CD38 upregulation following treatment to promote immunotherapy resistance. Additionally, the increased basal levels of Th17 cells in PD-L1^low^ 393P tumors suggests that Th17 cells may be a mechanism for primary immune evasion and tumor formation.

Although Th17 cells are commonly associated with a pro-inflammatory response^18^, analysis of RNA-seq data from melanoma patient biopsies during pre- and on-treatment of PD-1 blockade showed that a net increase in Th17-associated gene signatures correlated with poorer response and predicted poorer overall survival following treatment. These clinical findings corroborate our experimental data and identify Th17 genes as potential markers of resistance.

Our identification that the Th17-associated cytokine IL-17 promotes resistance to anti-PD-L1 alone or in combination with MEK inhibition validates a novel triple combinatorial therapeutic strategy for lung cancer patients, consistent with observations that IL-17 is involved in tumor progression^17, 21^. Moreover, the increase in cancer cell IL-6 expression from combination therapy as well as the role of Th17 in autoimmunity suggests that IL-17 blockade could potentially ameliorate the observed clinical adverse events due to immune checkpoint blockade^12, 34, 35, 36^. A major advantage of our proposed triple combinatorial treatment strategy is the fact that all three targets currently have drugs that have received FDA approval, as IL-17 drugs are FDA approved to treat psoriasis^37^. Moreover, results from our study emphasize the need for accessible lung cancer patient datasets with matched pre- and post-treatment biopsies to predict patient response and identify potential combination therapeutic strategies to improve patient survival.

## Materials and Methods

### Cell culture and reagents

Human and murine cell lines were cultured in RPMI 1640 (Gibco) with 10% fetal bovine serum (Gibco). Murine cell lines (Kras ^LA1/+^/p53^R172HΔg/+^) were derived as previously described^6^. Human cell lines were obtained from the American Type Culture Collection. All cell lines were routinely tested for mycoplasma contamination on a monthly basis using the LookOut Mycoplasma PCR Detection Kit (Sigma). Any cells yielding positive mycoplasma results were immediately discarded.

### Tumor models and in vivo treatments

Syngeneic tumor assays for male or female wild-type 129/sv mice were used. Cancer cells were injected subcutaneously into the mouse flank (unless otherwise noted 1 × 10^6^ cells in 100 μl of DPBS). Tumor size was calculated using the formula ½ (length x width^2^) at time point indicated. Lung metastatic nodules were counted after removal of lungs and counted as previously described^15^. Treatment was started at week 3 or when tumors were an average of 150-200 mm^3^ as measured by a digital caliper. Mice were randomized before receiving vehicle control, treatment, or combination therapy. Mice were treated with antibodies 200 μg of anti-PD-L1 (clone 10F.9G2), anti-PD1 (clone RMP1-14), or anti-IL17A (clone 17F3) per mouse or their IgG controls via i.p. injection once per week for the indicated time. Selumetinib (SelleckChem) (4% DMSO, 30% PEG300, 5% Tween80) or vehicle control was administered daily by oral gavage at 25 mg/kg mouse weight. Tumor sizes were measured weekly after treatment. All animal experiments were approved by the Institutional Animal Care and Use Committee at The University of Texas MD Anderson Cancer Center.

The Kras^LSL-G12D/+^; p53^fl/fl^ (KP^-/-^) adeno-Cre inducible mouse model (129/sv background) of lung adenocarcinoma was used per^38^. Male or female mice were infected with virus by intratracheal intubation at around 3 months of age. Drug treatment of these mice with Selumetinib and/or anti-PD-L1 antibody was started at 3 months post infection when the presence of tumors was confirmed by micro-CT imaging. Transverse cross-sectional CT images were analyzed for largest tumor area quantification. Tumor diameters were measured using image-J.

### Co-culture Assay

Spleens were isolated from tumor free male or female 129/sv mice of at least 3 months of age. Spleen was placed on a 0.45 μM strainer and a plunger was used to press the spleen through the strainer. Splenocytes collected in the flow through were further washed with FACS buffer and RBC lysis was performed following manufacturer recommendations (Biolegend). Cell count and viability check using trypan blue was performed before culturing splenocytes with tumor cells. Viable splenocytes were co-cultured with the indicated tumor cell line at a ratio of 1:20 (tumor cell line: splenocyte) in the presence of anti-CD3/ anti-CD28 (5 μg/ml) (Life technologies) for 96 hours at 37°C, 5% CO2. Where indicated AZD6244 (10 µM) and/or anti-PDL1 (20 µg/ml) were added at the time of cell seeding.

### Invasion/migration Assays

Transwell migration of 8µM inserts (BD-Biosciences) and invasion (BD-Bioscience) assays were performed for 16 hours as per^6^. Inserts were imaged using crystal violet solution and migratory or invasive cells analyzed on an Olympus IX73 microscope and counted using Image-J software. Five microscopy fields were taken per transwell insert for quantitation.

### Flow Cytometry Analysis

Tumors were processed following the MACS (Miltenyi) mouse tumor dissociation kit. Mechanical dissociation using a gentleMACS Octo dissociator (Miltenyi) was performed followed with enzymatic digestion with collagenase I (0.05% w/v, Sigma), DNase type IV (30 U/ml, Sigma), and hyaluronidase type V (0.01% w/v, Sigma) for 40 minutes with rocking at 37°C. Tumor samples were mechanically dissociated again and passed through a 70 μm filter before being stained with fluorochrome-conjugated antibodies in FACS buffer. RBC lysis (Biolegend) was performed on both single cell tumor and splenocytes samples following manufacturer recommendation. Cells were stained for surface markers using fluorochrome-conjugated anti-mouse antibodies (CD45, CD3, CD8, CD4) for 1 hour at room temperature. Ghost aqua BV510 (Tonobo) was used to stain dead cells. Cells were fixed using 1% PFA at room temperature for 15 minutes, then washed twice with perm/wash buffer (Biolegend). Cells were stained for intracellular antibodies at room temperature for 1 hour. Cells were filtered and analyzed on either a BD Canto II (BD Biosciences, San Jose CA), or BD LSR Fortessa (BD Biosciences) and analyzed using FlowJo software (v.10.5.3; Tree Star). For single color compensation, ultracomp eBeads compensation beads (Thermo Fisher) were used and stained with a single fluorescent-conjugated antibody according to manufacturer’s instructions. Compensation was calculated automatically using BD FACSDiva 8.0.1. The list and dilution of antibodies used is found in the Supplementary Methods.

### Protein Isolation and Western Blot analysis

Total cell lysates were obtained from human or murine cell lines following^7^. Protein was extracted from flash frozen tumors using a homogenizer (Fisher Scientific) in lysis buffer (1% Triton X-100, 50 mM HEPES, 150 mM NaCl, 1.5 mM MgCl2) on ice. The samples were pulsed for 5 seconds, until tissue was disintegrated and then incubated on ice for 30 minutes. 20 μg of lysate was loaded onto a SDS-PAGE gel, electrophoresed and transferred to a nitrocellulose membrane. After blocking with 5% fat free dry milk (Biorad) membranes were probed with primary antibodies listed in Table 1 overnight at 4°C. HRP-conjugated secondary antibodies were added to membranes and incubated for 1 hour. The Pierce ECL western blotting substrate (Thermo Fisher) was used. The protocol that was followed for all immunoblots is found in the following reference^7^.

### RNA isolation and Real-time qPCR analysis

RNA from tumors was isolated using the mirVana isolation kit (Thermo Fisher) per manufacturer recommendations. RNA from cell lines was isolated using TRIzol reagent (Thermo Fisher). After quantification of RNA using an Epoch plate reader Take3 plate 260/280nm (BioTek), 2 μg of RNA was used for reverse transcription using qSCRIPT master mix (Quanta biosciences). QPCR were performed using SYBR green PCR master mix (Thermo Fisher). Primers specific for each gene measured are listed in Table 2. L32 was used to normalize expression across all samples. QPCR reactions were performed on the 7500 Fast Real Time PCR system (Applied Biosystems), CFX96 Real-Time PCR detection system (Bio-Rad Laboratories), or CFX384 Real-Time PCR detection system (Bio-Rad Laboratories).

### Reverse phase protein array (RPPA)

Tumor tissues were mechanically homogenized in lysis buffer (1% Triton X-100, 50 mM HEPES [pH 7.4], 150 mM NaCl, 1.5 mM MgCl2, 1 mM EGTA, 100 mM NaF, 10 mM NaPPi, 10% glycerol, 1 mM phenylmethylsulfonyl fluoride, 1 mM Na3VO4, with protease and phosphoprotease inhibitors from Roche), incubated on ice for 20 minutes, centrifuged at 14,000 rpm for 10 minutes, and collected for supernatant. Protein concentration was measured using the Pierce BCA Protein Assay Kit (Thermo Fisher Scientific), and protein samples were prepared to a final concentration of 1 µg/µl after mixing with 4x SDS sample buffer (40% glycerol, 8% SDS, 0.25M Tris-HCl pH 6.8, 10% 2-mercaptoethanol) to produce a 1x SDS sample buffer solution. Protein samples were then boiled at 100°C for 5 minutes and stored at −80 °C for RPPA processing described here (https://www.mdanderson.org/research/research-resources/core-facilities/functional-proteomics-rppa-core/rppa-process.html). For analysis of RPPA data, a linear mixed model was applied to compare protein expression on a protein-by-protein basis between epithelial and mesenchymal groups; the model includes cell line effects as a random effect factor. The resulting p-values were modeled by a Beta-Uniform Model (BUM). To identify protein markers differentially regulated in epithelial and mesenchymal phenotypes, we used a False Discovery Rate (FDR) of 0.05 as the cutoff. The heatmap was generated based on mean adjusted expression for each cell line. The Pearson correlation was used for distance matrix calculation, and Ward method was applied as linkage rule for the hierarchical clustering.

### NanoString

129/sv mice implanted with 344SQ or 393P tumors were treated with anti-PD-L1 (200 µg/mouse) once per week i.p. and/or selumetinib (10 mg/kg) for 14 days. Tumors were harvested and flash frozen, total RNA isolated using the mirVana isolation kit (Thermo Fisher). Gene expression analysis was done using a custom mouse tumor microenvironment panel. A total of 100 ng of total RNA in a final volume of 5 μl was mixed with capture probes and reporter probes tagged with a fluorescent barcode from the custom gene expression code set. Probes and targets were hybridized at 65°C for 12-24 hours. Hybridized samples were run on the NanoString nCounter preparation station following manufacturer recommendations. The samples were run on maximum scan resolution on the nCounter Digital Analyzer. Data was analyzed using nSolver Analysis Software. Additional statistical analysis on NanoString nCounter data was conducted as described in^16^, using R version 3.4.2 (R Core Team, 2016). Samples were scaled by the geometric mean of the positive spike-in RNA hybridization controls, as described in the nCounter data analysis manual (NanoString Technologies, Inc., 2011). The expression of each endogenous gene was then tested against the detected expression of all negative control genes using 1-sided, 2-sample t tests. Genes showing greater expression than the negative control genes with p < 0.001 were included in further analysis, while others were omitted. Housekeeping gene normalization was applied using the same geometric mean scaling, with a housekeeping gene set identified as most stable in a larger data set by the method of^39^. This set consisted of Alas1, Abcf1, Tbp, Ppia, and Tubb5. Differential expression analysis was conducted on the data after log2 transformation, comparing treatment vs. IgG using the Empirical Bayes method in limma^40,41^.

### Immunohistochemistry (IHC)

Formalin-fixed paraffin-embedded (FFPE) tumor or lung tissue were cut to 4 µm sections and IHC was performed as follows: 3% hydrogen peroxide for 10 minutes, 5% goat serum blocking for 1 hour, tissues probed with primary antibody listed in Table 1 overnight in 4°C, secondary streptavidin-conjugated secondary antibody was probed for 1 hour, incubation with biotinylated HRP for 30 minutes, 3,3’-diaminobenzidine reagent (Dako) was used for signal development (∼5 minutes). Slides were counter-stained with Harris Hematoxylin (Thermo Fisher) and further dehydrated and mounted. IHC was performed following procedure adapted from Peng *et al*^7^. Digital images of stained slides were acquired using an Aperio slide scanner AT2 (Leica Biosystems). The Aperio imagescope software version 12.3.3 was used for further analysis.

### Analysis of RNA sequencing data

FPKM RNA sequence data previously published from Riaz et al^27^ was downloaded from GEO (GSE GSE91061). Patient samples with 0 sequencing reads were excluded in the analysis. Survival analysis was performed using the Kaplan-Meier method and log-rank test was used to determine statistical significance. Comparison of gene expression values between response groups was performed using the Wilcoxon matched-pairs signed rank test.

### Statistics

All statistical analysis were performed using GraphPad Prism software version 8.0.0. Unless otherwise noted, a one-way ANOVA post-hoc Tukey test was used for multi-group comparisons while unpaired Student t-test (two-tailed) was performed for two-group comparisons. A *p*-value of less than 0.05 was considered statistically significant.

## Financial Support

This was supported by: NIH R37CA214609, CPRIT-MIRA RP160652-P3, Rexanna’s Foundation for Fighting Lung Cancer, and LUNGevity Foundation award to D.L.G. & L.A.B. The work was also supported by the generous philanthropic contributions to The University of Texas MD Anderson Lung Cancer Moon Shots Program.

## Conflict of Interest

D.L.G. declares advisory board work for Janssen, AstraZeneca, GlaxoSmithKline and Sanofi. D.L.G. receives research grant funding from AstraZeneca, Janssen, Astellas, Ribon Therapeutics and Takeda. L.A.B. declares consulting work for AstraZeneca, AbbVie, GenMab, BergenBio, Pharma Mar, SA. L.A.B. receives research grant funding from AbbVie, AstraZeneca, GenMab, Tolero Pharmaceuticals. All other authors declare that they have no conflict of interests.

## Authors’ Contributions

Conceptualization, D.H.P., B.L.R., and D.L.G.; Investigation, C.A.G., L.D., J.O.K., J.W.; Writing, B.L.R, D.H.P., D.L.G.; Resources, A.P., P.O.G., J.J.F, L.G, L.C., L.A.B.; Supervision, D.L.G.

## Acknowledgements

This study made use of the Research Histology, Pathology, and Imaging Core, supported by P30 CA16672 DHHS/NCI Cancer Center Support Grant (CCSG). The Flow Cytometry Lab South Campus Core facility at MDACC is supported by CCSG NCI#P30 CA16672.

## References

1. Cox AD, Fesik SW, Kimmelman AC, Luo J, Der CJ. Drugging the undruggable RAS: Mission possible? Nat Rev Drug Discov 13, 828–851 (2014).

2. Wang Y, Kaiser CE, Frett B, Li H-y. Targeting Mutant KRAS for Anticancer Therapeutics: A Review of Novel Small Molecule Modulators. Journal of Medicinal Chemistry 56, 5219–5230 (2013).

3. Blumenschein GR, Jr., et al. A randomized phase II study of the MEK1/MEK2 inhibitor trametinib (GSK1120212) compared with docetaxel in KRAS-mutant advanced non-small-cell lung cancer (NSCLC)dagger. Ann Oncol 26, 894–901 (2015).

4. Jänne PA, et al. Selumetinib Plus Docetaxel Compared With Docetaxel Alone and Progression-Free Survival in Patients With KRAS-Mutant Advanced Non–Small Cell Lung Cancer. Jama 317, 1844 (2017).

5. Chen Z, et al. A murine lung cancer co-clinical trial identifies genetic modifiers of therapeutic response. Nature 483, 613–617 (2012).

6. Gibbons DL, et al. Contextual extracellular cues promote tumor cell EMT and metastasis by regulating miR-200 family expression. Genes Dev 23, 2140–2151 (2009).

7. Peng DH, et al. ZEB1 suppression sensitizes KRAS mutant cancers to MEK inhibition by an IL17RD-dependent mechanism. Sci Transl Med 11, (2019).

8. Chen L, et al. Metastasis is regulated via microRNA-200/ZEB1 axis control of tumour cell PD-L1 expression and intratumoral immunosuppression. Nat Commun 5, 5241 (2014).

9. Mak MP, et al. A Patient-Derived, Pan-Cancer EMT Signature Identifies Global Molecular Alterations and Immune Target Enrichment Following Epithelial-to-Mesenchymal Transition. Clin Cancer Res 22, 609–620 (2016).

10. Lou Y, et al. Epithelial-Mesenchymal Transition Is Associated with a Distinct Tumor Microenvironment Including Elevation of Inflammatory Signals and Multiple Immune Checkpoints in Lung Adenocarcinoma. Clin Cancer Res 22, 3630–3642 (2016).

11. Brahmer J, et al. Nivolumab versus Docetaxel in Advanced Squamous-Cell Non-Small-Cell Lung Cancer. N Engl J Med 373, 123–135 (2015).

12. Dong C. TH17 cells in development: an updated view of their molecular identity and genetic programming. Nat Rev Immunol 8, 337–348 (2008).

13. Cardnell RJ, et al. An Integrated Molecular Analysis of Lung Adenocarcinomas Identifies Potential Therapeutic Targets among TTF1-Negative Tumors, Including DNA Repair Proteins and Nrf2. Clin Cancer Res 21, 3480–3491 (2015).

14. Hennessy BT, et al. A Technical Assessment of the Utility of Reverse Phase Protein Arrays for the Study of the Functional Proteome in Non-microdissected Human Breast Cancers. Clin Proteomics 6, 129–151 (2010).

15. Chen L, et al. Metastasis is regulated via microRNA-200/ZEB1 axis control of tumour cell PD-L1 expression and intratumoral immunosuppression. Nat Commun 5, 5241 (2014).

16. Chen L, et al. CD38-Mediated Immunosuppression as a Mechanism of Tumor Cell Escape from PD-1/PD-L1 Blockade. Cancer Discov 8, 1156–1175 (2018).

17. Akbay EA, et al. Interleukin-17A Promotes Lung Tumor Progression through Neutrophil Attraction to Tumor Sites and Mediating Resistance to PD-1 Blockade. Journal of Thoracic Oncology 12, 1268–1279 (2017).

18. Bailey SR, Nelson MH, Himes RA, Li Z, Mehrotra S, Paulos CM. Th17 cells in cancer: the ultimate identity crisis. Front Immunol 5, 276 (2014).

19. Celada LJ, et al. PD-1 up-regulation on CD4(+) T cells promotes pulmonary fibrosis through STAT3-mediated IL-17A and TGF-beta 1 production. Science Translational Medicine 10, (2018).

20. Fabre T, et al. Type 3 cytokines IL-17A and IL-22 drive TGF-beta-dependent liver fibrosis. Sci Immunol 3, (2018).

21. Jin C, et al. Commensal Microbiota Promote Lung Cancer Development via gammadelta T Cells. Cell 176, 998–1013 e1016 (2019).

22. Korn T, Bettelli E, Oukka M, Kuchroo VK. IL-17 and Th17 Cells. Annu Rev Immunol 27, 485–517 (2009).

23. Liang SC, et al. Interleukin (IL)-22 and IL-17 are coexpressed by Th17 cells and cooperatively enhance expression of antimicrobial peptides. J Exp Med 203, 2271–2279 (2006).

24. Muranski P, Restifo NP. Essentials of Th17 cell commitment and plasticity. Blood 121, 2402–2414 (2013).

25. Punt S, Langenhoff JM, Putter H, Fleuren GJ, Gorter A, Jordanova ES. The correlations between IL-17 vs. Th17 cells and cancer patient survival: a systematic review. Oncoimmunology 4, e984547 (2015).

26. Hatton RD. TGF-beta in Th17 cell development: the truth is out there. Immunity 34, 288–290 (2011).

27. Riaz N, et al. Tumor and Microenvironment Evolution during Immunotherapy with Nivolumab. Cell 171, 934–949 e916 (2017).

28. Hellmann MD, et al. Phase Ib study of atezolizumab combined with cobimetinib in patients with solid tumors. Ann Oncol 30, 1134–1142 (2019).

29. Sullivan RJ, et al. Atezolizumab plus cobimetinib and vemurafenib in BRAF-mutated melanoma patients. Nat Med 25, 929–935 (2019).

30. Lee JW, et al. The Combination of MEK Inhibitor With Immunomodulatory Antibodies Targeting Programmed Death 1 and Programmed Death Ligand 1 Results in Prolonged Survival in Kras/p53-Driven Lung Cancer. J Thorac Oncol 14, 1046–1060 (2019).

31. Loi S, et al. RAS/MAPK Activation Is Associated with Reduced Tumor-Infiltrating Lymphocytes in Triple-Negative Breast Cancer: Therapeutic Cooperation Between MEK and PD-1/PD-L1 Immune Checkpoint Inhibitors. Clin Cancer Res 22, 1499–1509 (2016).

32. Ebert PJR, et al. MAP Kinase Inhibition Promotes T Cell and Anti-tumor Activity in Combination with PD-L1 Checkpoint Blockade. Immunity 44, 609–621 (2016).

33. Della Corte CM, et al. Antitumor activity of dual blockade of PD-L1 and MEK in NSCLC patients derived three-dimensional spheroid cultures. J Exp Clin Cancer Res 38, 253 (2019).

34. Callahan MK, et al. Evaluation of serum IL-17 levels during ipilimumab therapy: Correlation with colitis. Journal of Clinical Oncology 29, 2505–2505 (2011).

35. Uemura M, et al. Selective inhibition of autoimmune exacerbation while preserving the anti-tumor clinical benefit using IL-6 blockade in a patient with advanced melanoma and Crohn’s disease: a case report. J Hematol Oncol 9, 81 (2016).

36. Kimura A, Kishimoto T. IL-6: regulator of Treg/Th17 balance. Eur J Immunol 40, 1830–1835 (2010).

37. Campa M, Mansouri B, Warren R, Menter A. A Review of Biologic Therapies Targeting IL-23 and IL-17 for Use in Moderate-to-Severe Plaque Psoriasis. Dermatol Ther (Heidelb) 6, 1–12 (2016).

38. DuPage M, Dooley AL, Jacks T. Conditional mouse lung cancer models using adenoviral or lentiviral delivery of Cre recombinase. Nat Protoc 4, 1064–1072 (2009).

39. Vandesompele J, et al. Accurate normalization of real-time quantitative RT-PCR data by geometric averaging of multiple internal control genes. Genome Biol 3, RESEARCH0034 (2002).

40. Ritchie ME, et al. limma powers differential expression analyses for RNA-sequencing and microarray studies. Nucleic Acids Res 43, e47 (2015).

41. Smyth GK. Linear models and empirical bayes methods for assessing differential expression in microarray experiments. Stat Appl Genet Mol Biol 3, Article3 (2004).

