## Supplemental Materials for "Th17 cells contribute to combination MEK inhibitor and anti-PD-L1 therapy resistance in *KRAS/p53* mutant lung cancers"

Supplementary Figure 1

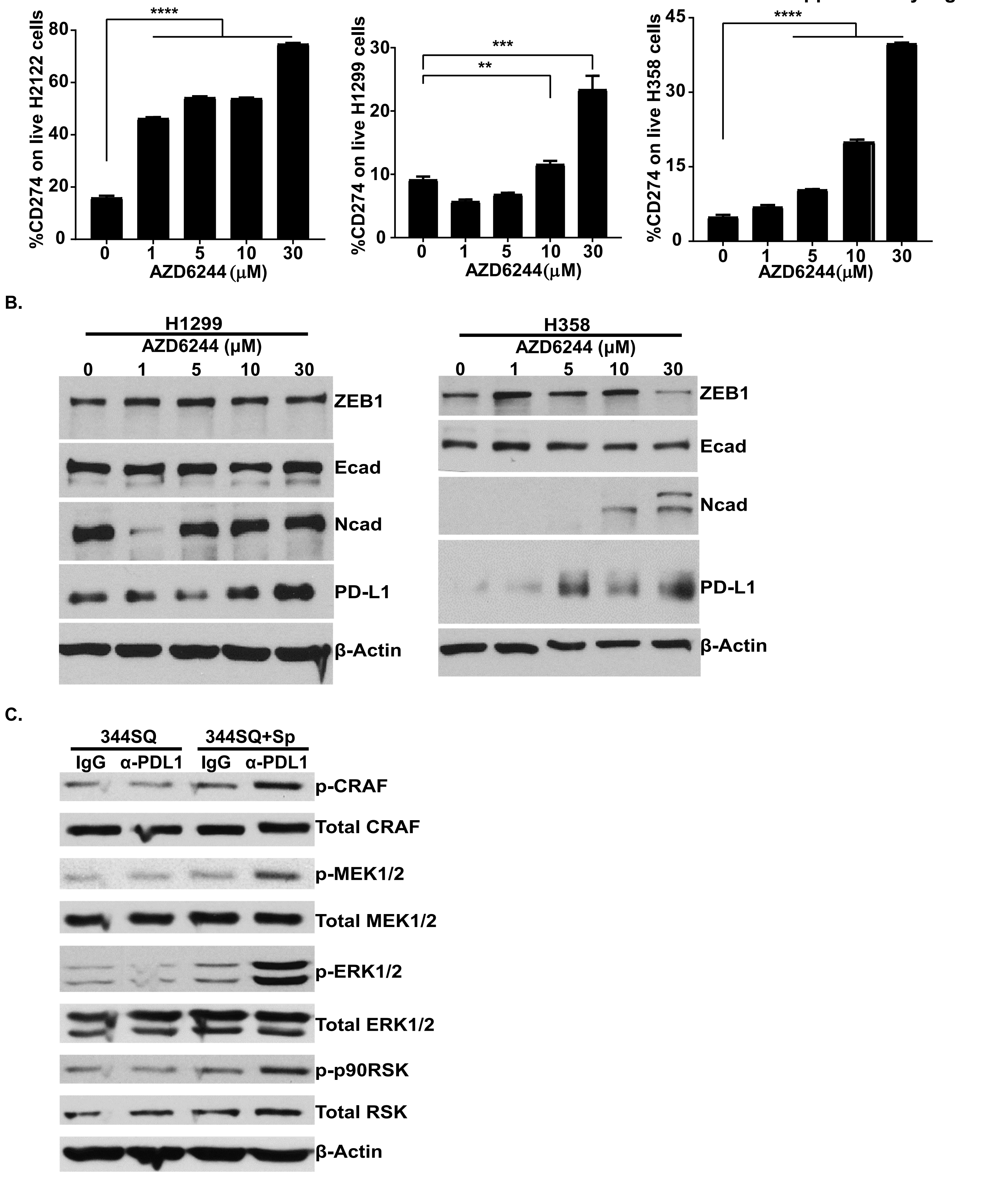

**Supplementary Figure Legends**

**Supplementary Figure S1.**

**(A)** Percent CD274+ (PD-L1) human lung cancer cell lines following 48 hr treatment with indicated concentrations of AZD6244.

**(B)** Western blot of indicated proteins in H358 and H1299 human lung cancer cell lines following 48 hr treatment with indicated concentrations of AZD6244.

**(C)** Western blot of indicated proteins in 344SQ murine KP cell line cultured alone or in co-culture with splenocytes, treated with 20 μg/ml anti-PD-L1 blocking antibody for 96 hrs.

Supplementary Figure 2

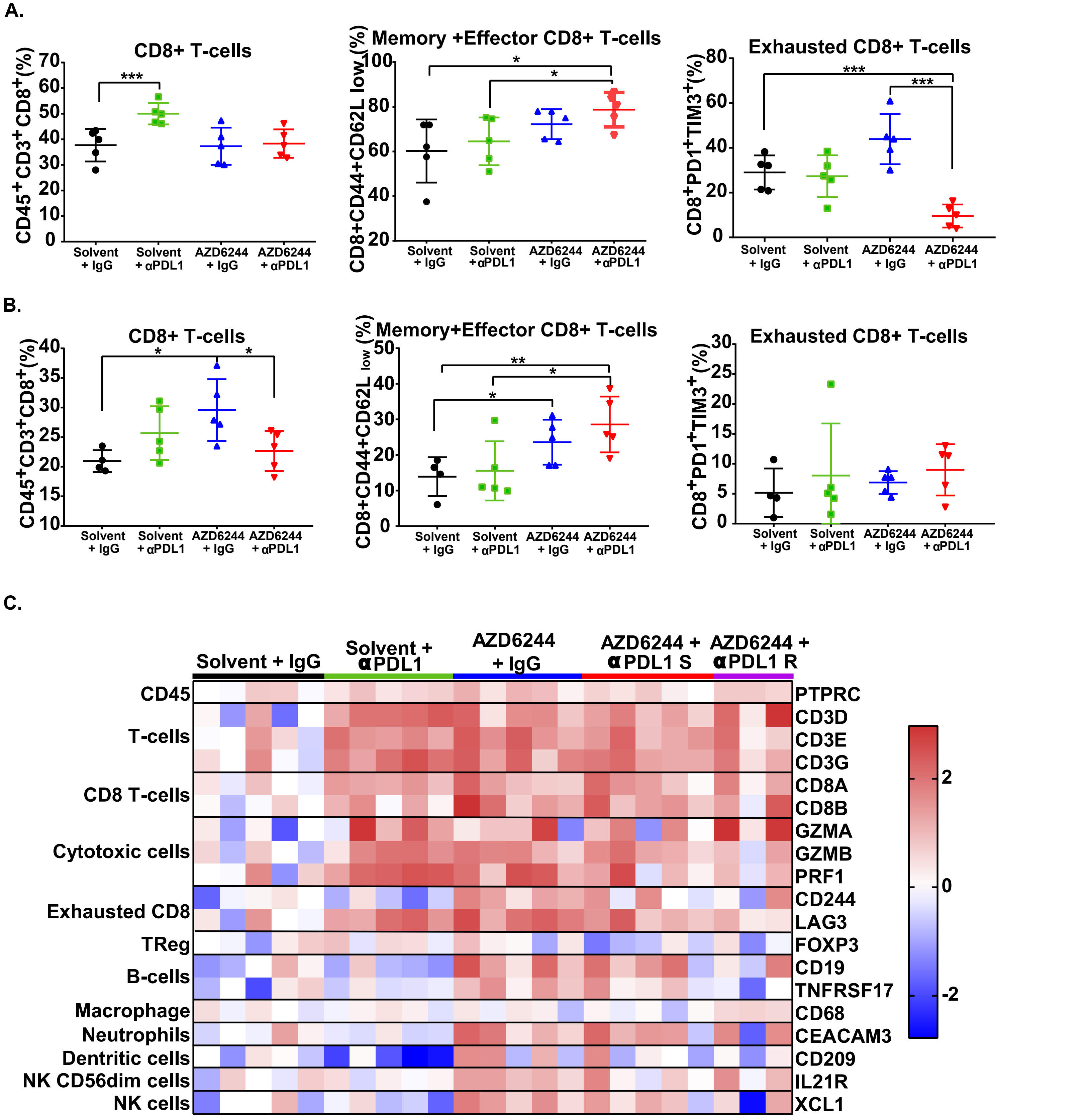

Supplementary Figure 2

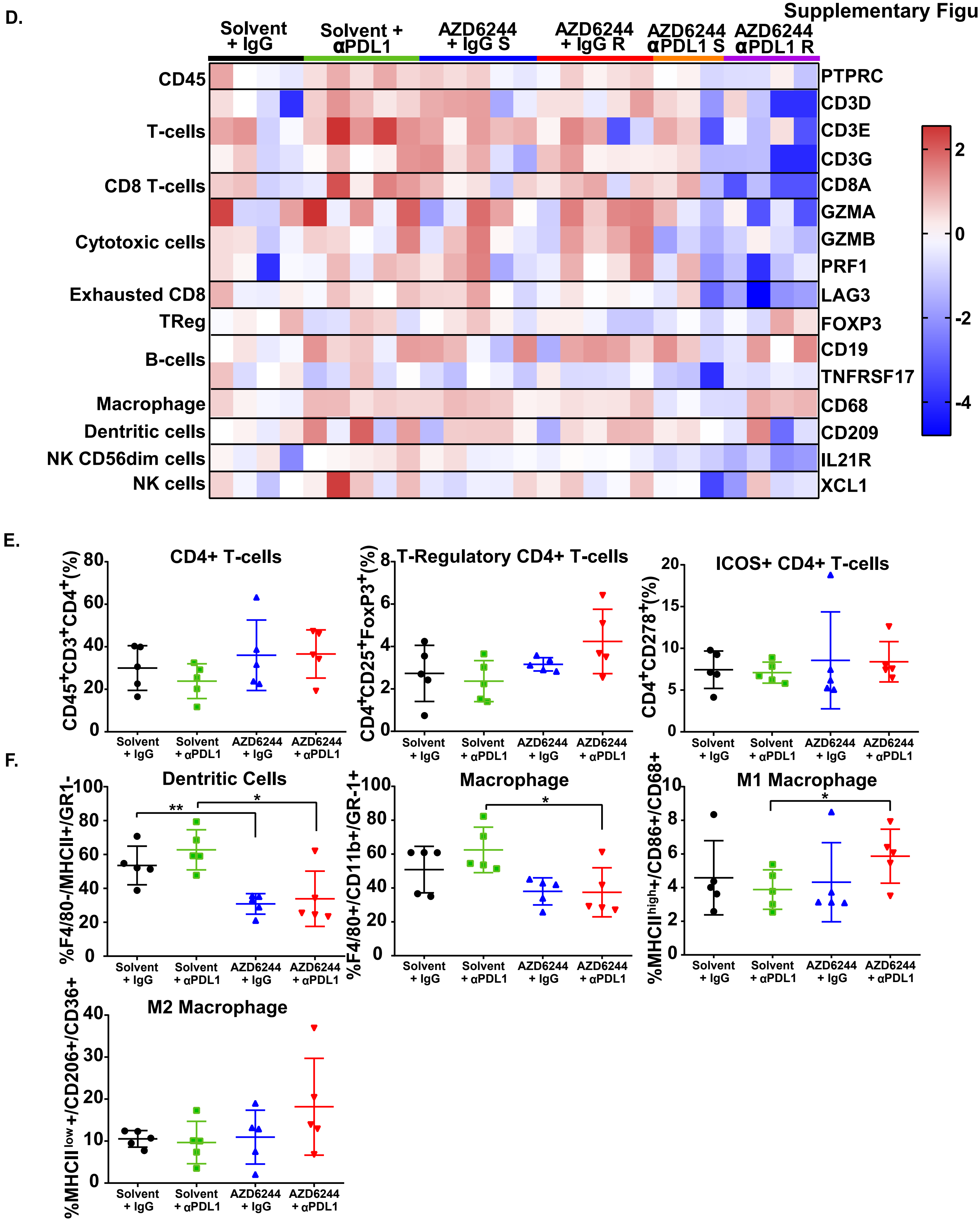

G.

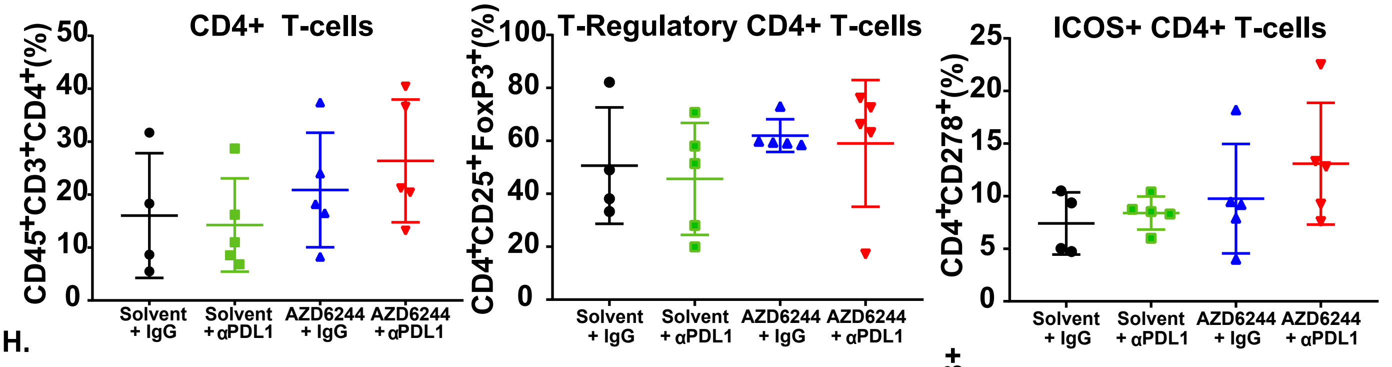
CD4+ T-cells
CD45+CD3+CD4+(%)
T-Regulatory CD4+ T-cells
CD4+CD25+FoxP3+(%)
ICOS+ CD4+ T-cells
CD4+CD278+(%)
Solvent + IgG
Solvent + αPDL1
AZD6244 + IgG
AZD6244 + αPDL1

H.

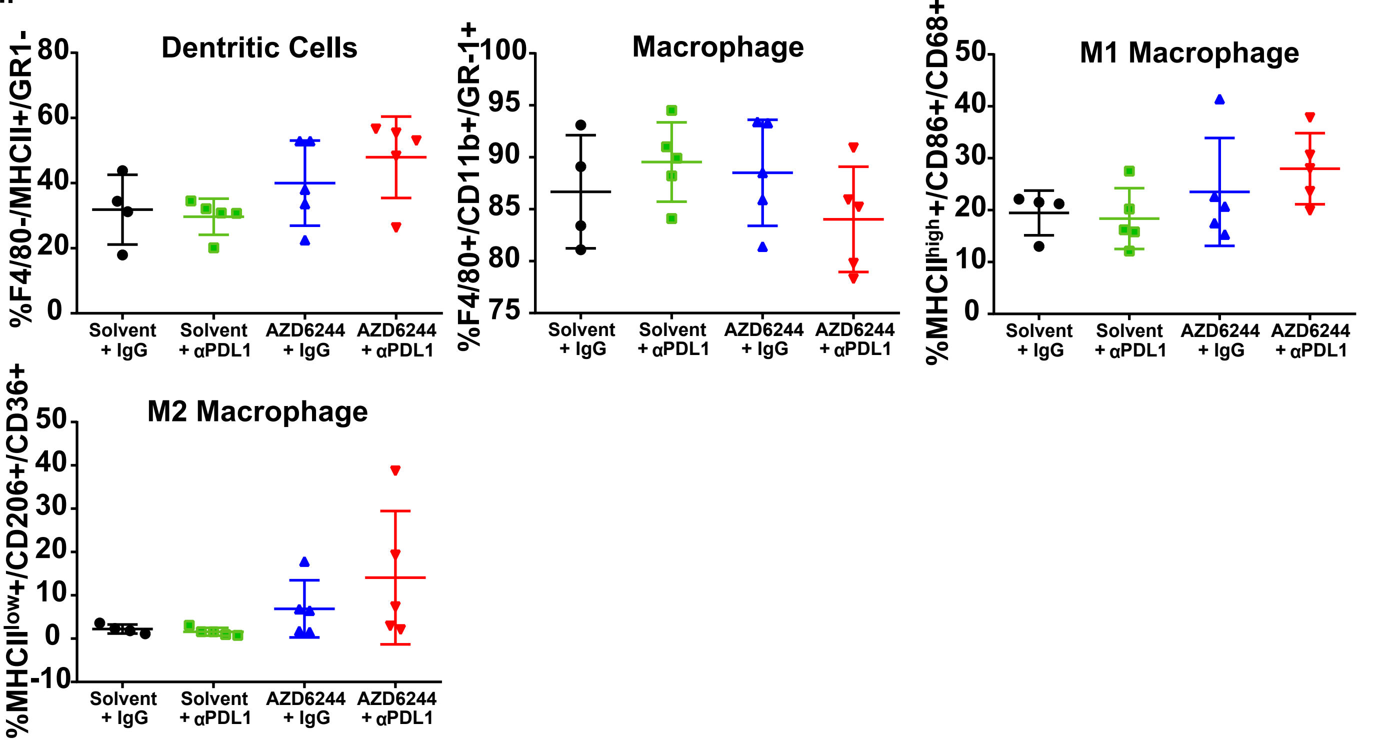
Dentritic Cells
%F4/80-/MHCII+/GR1-
Macrophage
%F4/80+/CD11b+/GR-1+
M1 Macrophage
%MHCIIhigh+/CD86+/CD68+
M2 Macrophage
%MHCIIlow+/CD206+/CD36+
Solvent + IgG
Solvent + αPDL1
AZD6244 + IgG
AZD6244 + αPDL1

**Supplementary Figure S2.**

**(A)** Percentage of total CD8+ T cells gated from CD45+CD3+ cells and CD8+ T cell subpopulations gated from total CD8+ T cells in 344SQ tumors treated with 25 mg/kg AZD6244 daily and/or weekly PD-L1 blocking antibody alone or in combination for 2 weeks.

**(B)** Percentage of total CD8+ T cells gated from CD45+CD3+ cells and CD8+ T cell subpopulations gated from total CD8+ T cells using indicated markers in 393P tumors treated with 25 mg/kg AZD6244 daily and/or weekly PD-L1 blocking antibody alone or in combination for 2 weeks.

**(C)** Heatmap of nanostring analysis of immune-related gene expression from 344SQ tumor tissues with indicated treatment groups at experimental endpoint from Figure 2A.

**(D)** Heatmap of nanostring analysis of immune-related gene expression from 344SQ tumor tissues with indicated treatment groups at experimental endpoint from Figure 2D.

**(E)** Percentage of total CD4+ T cells gated from CD45+CD3+ cells and CD4+ T cell subpopulations gated from total CD4+ T cells in 344SQ tumors treated with 25 mg/kg AZD6244 daily and/or weekly PD-L1 blocking antibody alone or in combination for 2 weeks.

**(F)** Percentage of indicated antigen presenting cell (APC) populations gated from CD45+ cells in 344SQ tumors treated with 25 mg/kg AZD6244 daily and/or weekly PD-L1 blocking antibody alone or in combination for 2 weeks.

**(G)** Percentage of total CD4+ T cells gated from CD45+CD3+ cells and CD4+ T cell subpopulations gated from total CD4+ T cells in 393P tumors treated with 25 mg/kg AZD6244 daily and/or weekly PD-L1 blocking antibody alone or in combination for 2 weeks.

**(H)** Percentage of indicated antigen presenting cell (APC) populations gated from CD45+ cells in 393P tumors treated with 25 mg/kg AZD6244 daily and/or weekly PD-L1 blocking antibody alone or in combination for 2 weeks.

Supplementary Figure 3

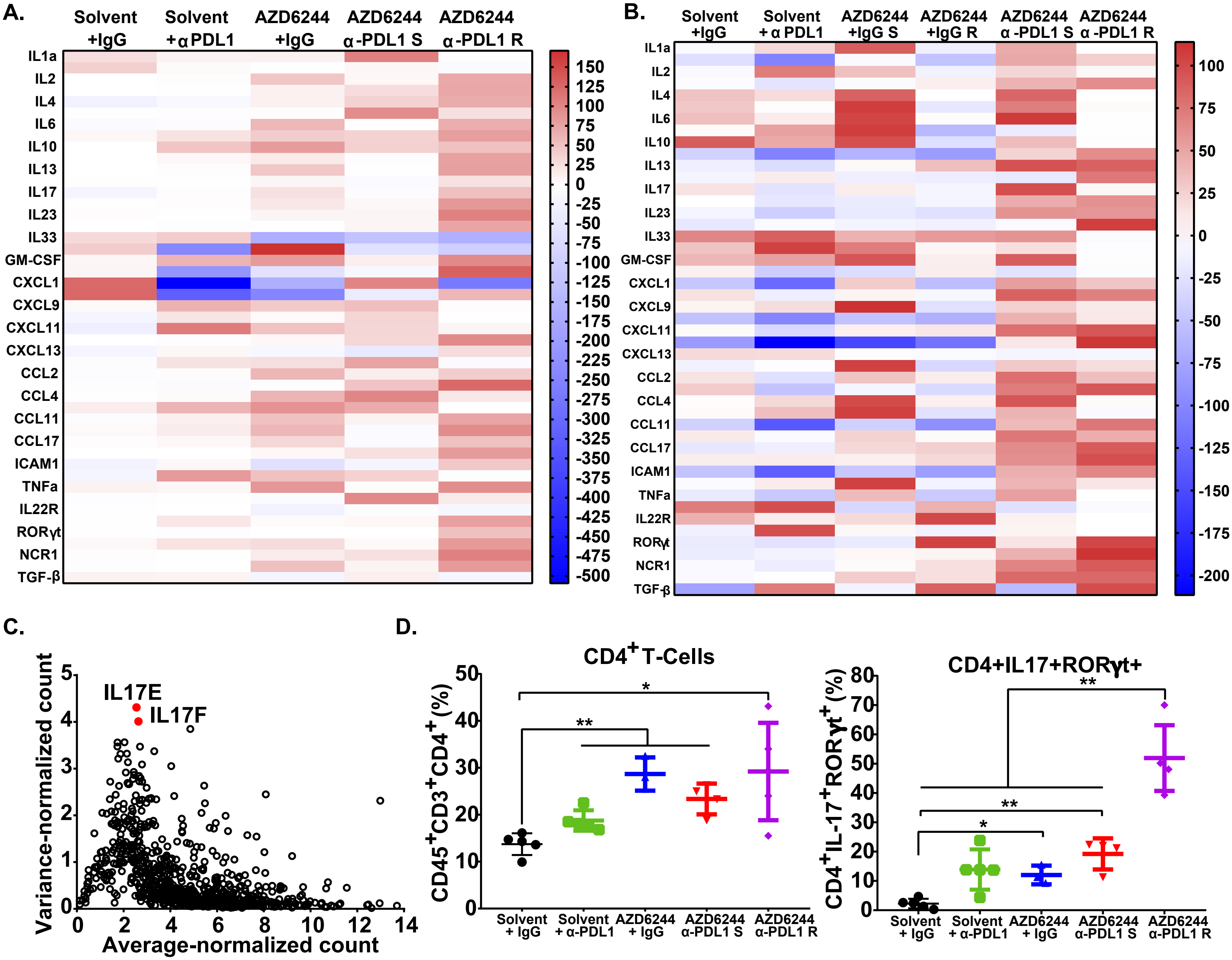

**Supplementary Figure S3.**

**(A)** Cytokine qPCR array heatmap of 344SQ tumors from the experiment in Figure 2A.

**(B)** Cytokine qPCR array heatmap of 393P tumors from the experiment in Figure 2D.

**(C)** Nanostring gene expression levels from 344SQ tumors treated with combinatorial AZD6244 and anti-PD-L1 therapies in Figure 2A at the point of sensitivity (Combo-S) versus point of resistance (Combo-R).

**(D)** Left: Percentage of total CD4+ T cells gated from CD45+CD3+ cells in 344SQ tumors with indicated treatments from the experiment in Figure 2A. Right: Percentage of IL-17+RORγt+ Th17 cells gated from total CD4+ T cells (left) in 344SQ tumors with indicated treatments from the experiment in Figure 2A.

Supplementary Figure 4

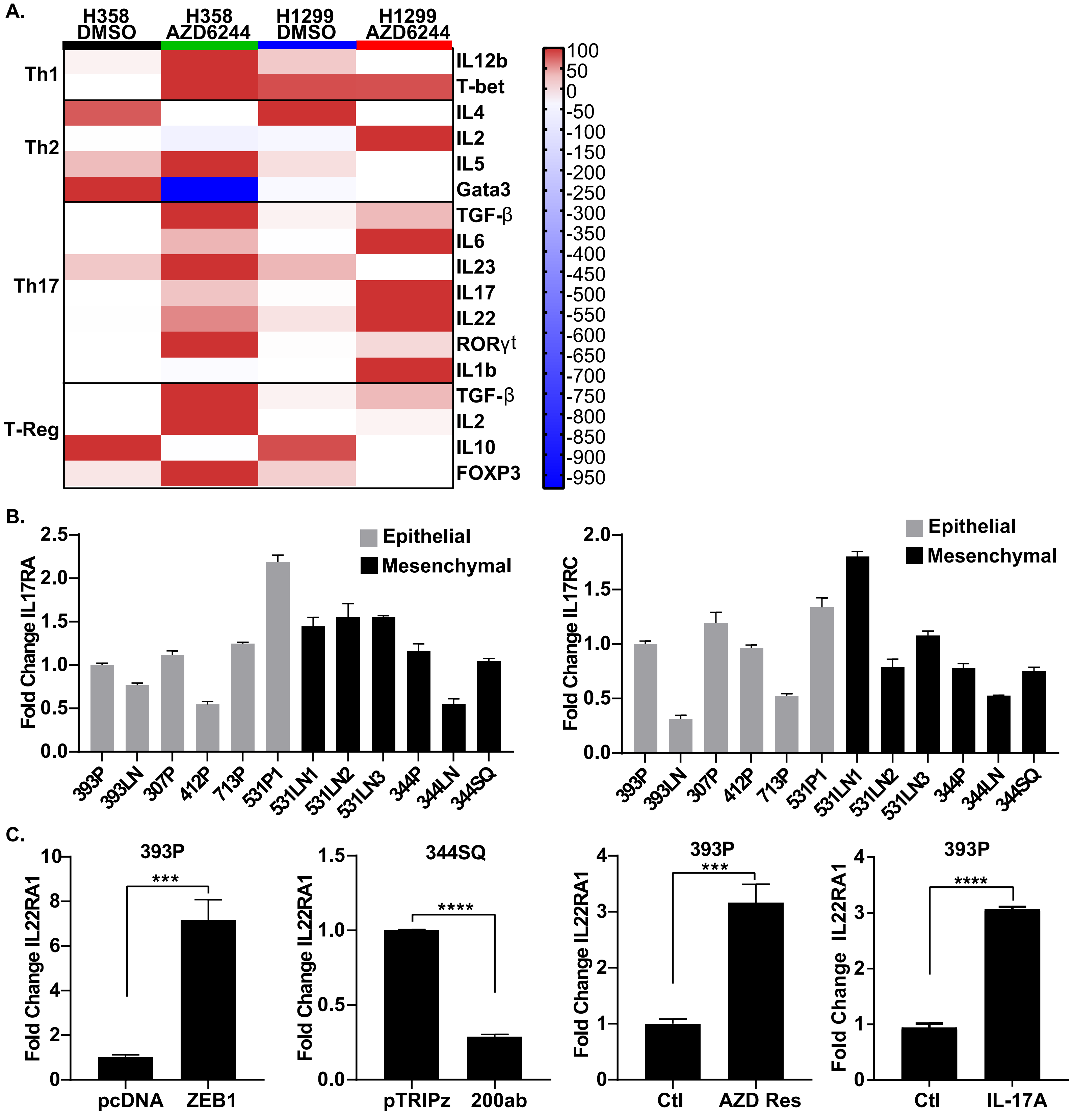

**Supplementary Figure S4.**

**(A)** Cytokine qPCR array heatmap of H358 and H1299 human lung cancer cell lines following treatment with DMSO control or 10 μM AZD6244 for 48 hrs.

**(B)** Left: QPCR analysis of IL17RA expression in panel of murine epithelial or mesenchymal cells.

Right: QPCR analysis of IL17RC expression in panel of murine epithelial or mesenchymal cells.

**(C)** QPCR analysis of IL22RA1 in 393P mouse KP cells with pcDNA vector control or constitutive ZEB1 expression, 344SQ mouse KP cells with pTRIPz vector control or miR200ab expression, parental 393P cells or 393P cells that developed AZD6244 resistance, and 393P cells treated with solvent control or recombinant mouse IL-17A.

Supplementary Figure 5

A.

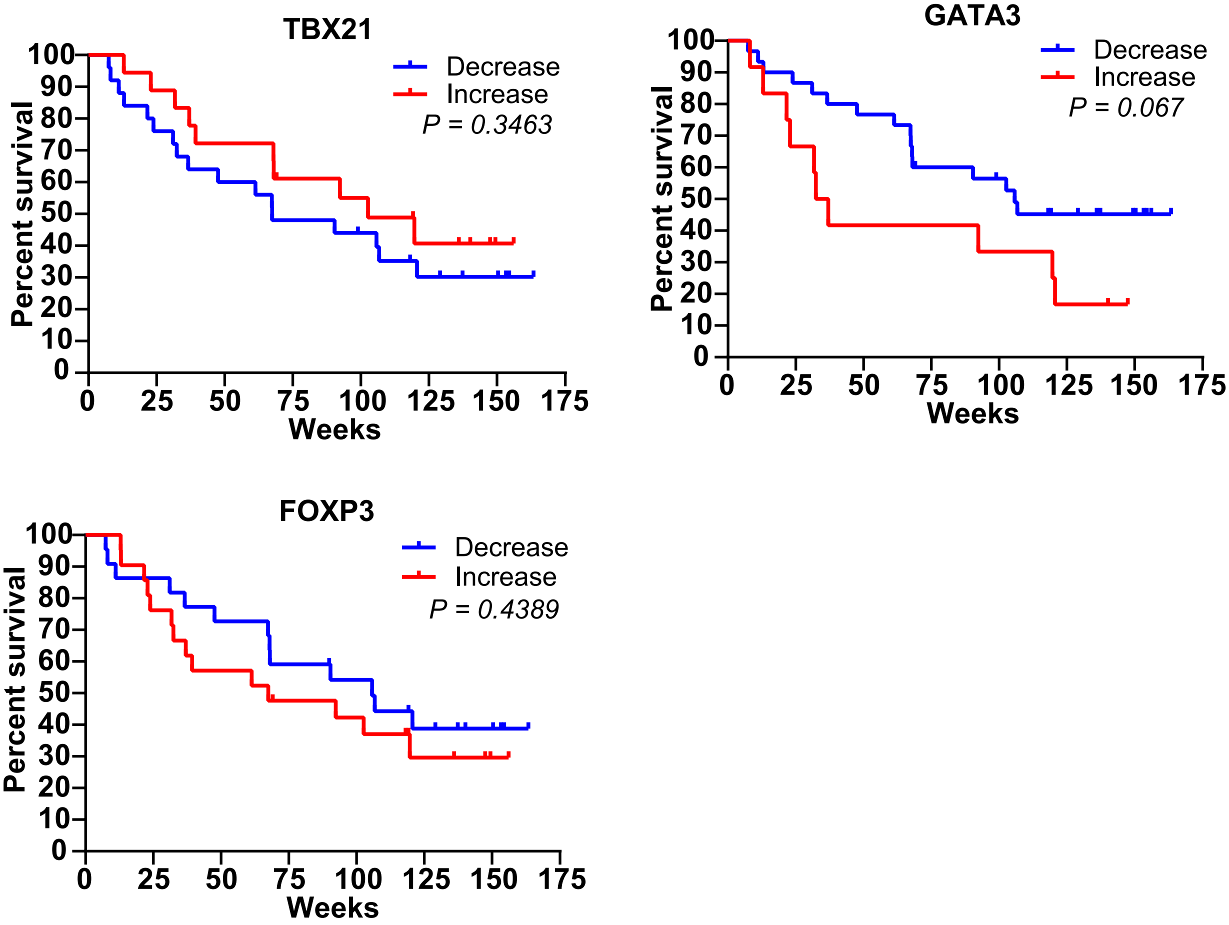


B.

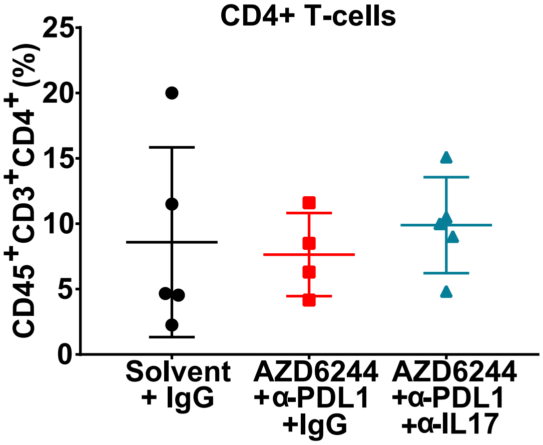

**Supplementary Figure S5.**

**(A)** Kaplan-Meier curves predicting survival of nivolumab patients based on changes of TBX21, GATA3, and FOXP3. Statistical differences was determined using Log-rank Cox test.

**(B)** Percent of CD45+CD3+CD4+ total T- cells from tumors treated with Solvent, AZD6244 + aPDL1, and AZD6244+a-PDL1+anti-IL17.

**Supplementary Methods**

QPCR Primers:

| | |
|---|---|
| mL32-F | GGAGAAGGTTCAAGGGCCAG |
| mL32-R | TGCTCCCATAACCGATGTGT |
| mCXCL13-F1 | CATAGATCGGATTCAAGTTACGCC |
| mCXCL13-R1 | GTAACCATTTGGCACGAGGATTC |
| mG-CSF-F1 | ATCCCGAAGGCTTCCCTGAGTG |
| mG-CSF-R1 | AGGAGACCTTGGTAGAGGCAGA |
| mGM-CSF-F1 | AACCTCCTGGATGACATGCCTG |
| mGM-CSF-R1 | AAATTGCCCCGTAGACCCTGCT |
| mCCL1-F1 | GCTTACGGTCTCCAATAGCTGC |
| mCCL1-R1 | GCTTTCTCTACCTTTGTTCAGCC |
| mCCL11-F1 | TCCATCCCAACTTCCTGCTGCT |
| mCCL11-R1 | CTCTTTGCCCAACCTGGTCTTG |
| mICAM1-F1 | AAACCAGACCCTGGAACTGCAC |
| mICAM1-R1 | GCCTGGCATTTCAGAGTCTGCT |
| mIFNg-F1 | CAGCAACAGCAAGGCGAAAAAGG |
| mIFNg-R1 | TTTCCGCTTCCTGAGGCTGGAT |
| mIL1a-F1 | ACGGCTGAGTTTCAGTGAGACC |
| mIL1a-R1 | CACTCTGGTAGGTGTAAGGTGC |
| mIL1b-F1 | TGGACCTTCCAGGATGAGGACA |
| mIL1b-R1 | GTTCATCTCGGAGCCTGTAGTG |
| mIL2-F1 | GCGGCATGTTCTGGATTTGACTC |
| mIL2-R1 | CCACCACAGTTGCTGACTCATC |
| mIL3-F1 | CCTGCCTACATCTGCGAATGAC |
| mIL3-R1 | GAGGTTAGCACTGTCTCCAGATC |
| mIL4-F1 | ATCATCGGCATTTTGAACGAGGTC |
| mIL4-R1 | ACCTTGGAAGCCCTACAGACGA |
| mIL5-F1 | GATGAGGCTTCCTGTCCCTACT |
| mIL5-R1 | TGACAGGTTTTGGAATAGCATTTCC |
| mIL7-F1 | CAGGAACTGATAGTAATTGCCCG |
| mIL7-R1 | CTTCAACTTGCGAGCAGCACGA |
| mIL10-F1 | CGGGAAGACAATAACTGCACCC |
| mIL10-R1 | CGGTTAGCAGTATGTTGTCCAGC |
| mIL12B-F1 | TTGAACTGGCGTTGGAAGCACG |
| mIL12B-R1 | CCACCTGTGAGTTCTTCAAAGGC |
| mIL16-F1 | CACGCAGACTTCATCCTCCACA |
| mIL16-R1 | AGCTATAGTCCATCCGTGCCTG |
| mIL27-F1 | TCTCGATTGCCAGGAGTGAACC |

| | |
|---|---|
| mIL27-R1 | AGTGTGGTAGCGAGGAAGCAGA |
| mCXCL1-F1 | TCCAGAGCTTGAAGGTGTTGCC |
| mCXCL1-R1 | AACCAAGGGAGCTTCAGGGTCA |
| mM-CSF-F1 | GCCTCCTGTTCTACAAGTGGAAG |
| mM-CSF-R1 | ACTGGCAGTTCCACCTGTCTGT |
| mCCL12-F1 | GCTACAGGAGAATCACAAGCAGC |
| mCCL12-R1 | ACGTCTTATCCAAGTGGTTTATGG |
| mCCL3-F1 | ACTGCCTGCTGCTTCTCCTACA |
| mCCL3-R1 | ATGACACCTGGCTGGGAGCAAA |
| mCCL4-F1 | ACCCTCCCACTTCCTGCTGTTT |
| mCCL4-R1 | CTGTCTGCCTCTTTTGGTCAGG |
| mCXCL12-F1 | CATCCAGAGCTTGAGTGTGACG |
| mCXCL12-R1 | GGCTTCAGGGTCAAGGCAAACT |
| mCCL5-F1 | CCTGCTGCTTTGCCTACCTCTC |
| mCCL5-R1 | ACACACTTGGCGGTTCCTTCGA |
| mCXCL12-F1 | GGAGGATAGATGTGCTCTGGAAC |
| mCXCL12-R1 | AGTGAGGATGGAGACCGTGGTG |
| mCCL17-F1 | CGAGAGTGCTGCCTGGATTACT |
| mCCL17-R1 | GGTCTGCACAGATGAGCTTGCC |
| mTNFa-F1 | GGTGCCTATGTCTCAGCCTCTT |
| mTNFa-R1 | GCCATAGAACTGATGAGAGGGAG |
| mIL17A-F1 | CAGACTACCTCAACCGTTCCAC |
| mIL17A-R1 | TCCAGCTTTCCCTCCGCATTGA |
| mIL22-F1 | GCTTGAGGTGTCCAACTTCCAG |
| mIL22-R1 | ACTCCTCGGAACAGTTTCTCCC |
| mIL21-F1 | GCCTCCTGATTAGACTTCGTCAC |
| mIL21-R1 | CAGGCAAAAGCTGCATGCTCAC |
| mIL23a-F1 | CATGCTAGCCTGGAACGCACAT |
| mIL23a-R1 | ACTGGCTGTTGTCCTTGAGTCC |
| mIL23R-F1 | GTCCACCAAACTTCCCAGACAG |
| mIL23R-R1 | CCTGAAGCAGGATGTCCTCTGA |
| mIL6-F1 | TACCACTTCACAAGTCGGAGGC |
| mIL6-R1 | CTGCAAGTGCATCATCGTTGTTC |
| mIL17RA-F1 | CTGTATGACCTGGAGGCTTTCTG |
| mIL17RA-R1 | CGAGTAGACGATCCAGACCTTC |
| mIL17RC-F1 | TAGAGCCAGACTCTGAGAGGGT |
| mIL17RC-R1 | AAGGCGCATCTAGCTGCCATAC |
| mIL22RA1-F1 | TTTCCTCGTCGGCTTGCTCTGT |
| mIL22RA1-R1 | CGTGTTCTTGGATGAAGCGTAGG |
| mTGFB1-F1 | TGATACGCCTGAGTGGCTGTCT |

| | |
|---|---|
| mTGFB1-R1 | CACAAGAGCAGTGAGCGCTGAA |
| mTbx21-F1 | CCACCTGTTGTGGTCCAAGTTC |
| mTbx21-R1 | CCACAAACATCCTGTAATGGCTTG |
| mIL25-F1 | TGGCTGAAGTGGAGCTCTGCAT |
| mIL25-R1 | CCCGATTCAAGTCCCTGTCCAA |

Antibodies:

| Antigen | Vendor | Catalog Number | Application |
|---|---|---|---|
| PD-L1 | Abcam | ab213480 | WB |
| β-actin | Sigma-Aldrich | A1978 | WB |
| Zeb1 | Santa Cruz | sc-25388 | WB |
| Mouse LOXL2 | Santa Cruz | sc-66950 (H-65) | WB |
| N-cadherin | BD Biosciences | 610920 | WB |
| E-cadherin | BD Biosciences | 610181 | WB |
| Vimentin (Western) | Cell Signaling | cs- 3932 | WB |
| p-CRaf (S338) | Cell Signaling | cs-9427 | WB |
| Total CRaf | BD Biosciences | 610151 | WB |
| p-Mek1/2 (S221/222) | Cell Signaling | cs-9154 | WB |
| Total Mek1/2 | Cell Signaling | cs-9122 | WB |
| p-Erk1/2 (T202/Y204) | Cell Signaling | cs-9101 | WB |
| Total Erk1/2 | Cell Signaling | cs-9102 | WB |
| p-p90RSK (Thr359/Ser363) | Cell Signaling | 9344S | WB |
| Total RSK1/2/3 | Cell Signaling | 9355S | WB |
| CD3 PE-594 | Biolegend | 100246 | FACS |
| CD4 APC-Cy7 | Biolegend | 100526 | FACS |
| CD8 PE-Cy7 | Biolegend | 100721 | FACS |
| CD44 BV711 | Biolegend | 103057 | FACS |
| CD62L FITC | Tonbo Biosciences | 35-0621-U500 | FACS |
| CD69 BV650 | Biolegend | 104541 | FACS |
| PD-1 BV605 | Biolegend | 135220 | FACS |
| TIM-3 APC | Biolegend | 134007 | FACS |
| LAIR1 PE | Invitrogen | 12-3051-82 | FACS |
| Live/Dead Ghost Violet 510 | Tonbo Biosciences | 13-0870-T100 | FACS |
| ICOS (CD278) BV786 | Biolegend | 313510 | FACS |
| CD25 BV395 | Biolegend | 564022 | FACS |
| FOXP3 PerCP-Cy5.5 | Invitrogen | 45-5773-82 | FACS |
| CD11b BV650 | Biolegend | 101239 | FACS |
| CD11c BV785 | Biolegend | 117335 | FACS |
| GR-1 BV711 | Biolegend | 108443 | FACS |
| F4/80 APC | Tonbo Biosciences | 20-4801-U100 | FACS |
| MHCII PE-Cy7 | Biolegend | 107629 | FACS |
| CD45 PerCP-Cy5.5 | Biolegend | 103132 | FACS |
| CCR5 FITC | Biolegend | 313705 | FACS |
| CCR6 BV421 | Biolegend | 129817 | FACS |

| | | | |
|---|---|---|---|
| RORγt PE and no color | ThermoFisher | 12-6988-82, 14-6988-82 | FACS and IHC |
| IL-17A APC | Biolegend | 506916 | FACS |
